# A nuclear architecture screen in *Drosophila* identifies Stonewall as a link between chromatin position at the nuclear periphery and germline stem cell fate

**DOI:** 10.1101/2023.11.17.567611

**Authors:** Ankita Chavan, Randi Isenhart, Son C. Nguyen, Noor Kotb, Jailynn Harke, Anna Sintsova, Gulay Ulukaya, Federico Uliana, Caroline Ashiono, Ulrike Kutay, Gianluca Pegoraro, Prashanth Rangan, Eric F. Joyce, Madhav Jagannathan

## Abstract

The association of genomic loci to the nuclear periphery is proposed to facilitate cell-type specific gene repression and influence cell fate decisions. However, the interplay between gene position and expression remains incompletely understood, in part because the proteins that position genomic loci at the nuclear periphery remain unidentified. Here, we used an Oligopaint-based HiDRO screen targeting ∼1000 genes to discover novel regulators of nuclear architecture in *Drosophila* cells. We identified the heterochromatin-associated protein, Stonewall (Stwl), as a factor promoting perinuclear chromatin positioning. In female germline stem cells (GSCs), Stwl binds and positions chromatin loci, including GSC differentiation genes, at the nuclear periphery. Strikingly, Stwl-dependent perinuclear positioning is associated with transcriptional repression, highlighting a likely mechanism for Stwl’s known role in GSC maintenance and ovary homeostasis. Thus, our study identifies perinuclear anchors in *Drosophila* and demonstrates the importance of gene repression at the nuclear periphery for cell fate.

## Introduction

The distribution of the genome within the interphase nucleus can tune cell-specific gene expression. In both plant and animal cells, dense-staining heterochromatin and repressed tissue-specific genes are typically found near the inner nuclear membrane (INM)^1^. In metazoans, an INM-associated network, involving the intermediate filament protein lamin and other associated proteins, serves as a scaffold for the organization of peripheral chromatin^2^. This chromatin, which is associated with the nuclear lamina, is referred to as lamina-associated domains (LADs) and is usually gene-poor, transcriptionally silent, and rich in repressive histone marks^3–6^. Experiments using LAD-embedded transcriptional reporters^4,7–9^ and gene tethering to the nuclear periphery^10–12^ have shown that perinuclear positioning is generally associated with reduced transcriptional output, although exceptions can occur^12^. Functionally, perinuclear positioning of a locus has been speculated to preserve the inactive transcriptional state and stabilize cell-specific gene expression programs^1,13^. Consistently, detachment of specific loci from the nuclear periphery in multiple cell types is associated with ectopic gene expression and alterations in cell fate decisions^14–17^. While the nuclear lamina^14–16,18^, nuclear pore complex (NPC) proteins^19–21^ and epigenetic modifications^22–24^ are known to influence chromatin association to the nuclear periphery, very few chromatin-binding perinuclear anchors have been identified thus far^17,25,26^. As a result, the precise relationships between perinuclear positioning, gene expression and cell fate remain enigmatic.

In this study, we leverage our recently developed HiDRO technology^27^ to conduct an RNAi screen in *Drosophila* cells aimed at identifying perinuclear anchors for heterochromatin. We individually depleted approximately 1,000 genes known to possess characteristic DNA-binding domains or nuclear localization sequences, and then measured changes in the spatial positioning of genomic regions located both at the periphery and center of the nucleus. Among our hits, we isolated a significant hit, the heterochromatin-associated MADF-BESS domain containing protein, Stonewall (Stwl)^28^ as a factor important for the peripheral positioning of LAD-enriched chromatin. MADF-BESS proteins are transcriptional regulators that bind DNA through an N-terminal MADF (Myb-SANT like in ADF) domain, whereas the C-terminal BESS motif mediates protein-protein interactions^29,30^. Previous studies have demonstrated that Stwl has a cell-autonomous function in female germline stem cell (GSC) maintenance^28,31,32^ as well as later stages of oogenesis^28,31,33,34^, likely through gene repression. Notably, Stwl-depleted GSCs are reported to differentiate precociously (as determined by fusome-containing germline cysts), even in the absence of critical differentiation genes^31^, suggesting that Stwl plays an important role in the balance between GSC self-renewal and differentiation. However, the mechanism by which Stwl fine-tunes this vital regulatory step in GSC cell fate has remained unclear. Here, we show that Stwl is crucial for perinuclear chromatin positioning in female GSCs. Using RNA sequencing, chromatin profiling and single molecule FISH, we demonstrate that Stwl promotes repression of canonical GSC differentiation genes such as *benign gonial cell neoplasm* (*bgcn*) by positioning these gene loci at the nuclear periphery. Overall, our HiDRO screen has identified multiple factors regulating nuclear architecture in *Drosophila*. In particular, we have pinpointed Stwl as an important factor that links perinuclear chromatin organization to female GSC fate.

## Results

### Discovery of novel regulators of chromosome positioning

To identify proteins involved in the positioning of chromatin at the nuclear periphery, we performed an RNAi screen using our recently developed HiDRO platform^27^ in Drosophila Kc167 cells (**Figure 1A**). Specifically, we seeded Kc167 cells onto 384-well plates containing individual dsRNAs in each well and performed high-throughput Oligopaint FISH to mark three 1Mb genomic regions that span Chromosome 2R and contain varying amounts of LADs (referred to as Chr. 2R-A, -B, and -C)^35^. In particular, 74% of Chr. 2R-C is designated as LADs in Kc167 cells (**Figure 1B**). We also confirmed by high-resolution FISH that this region was in closer proximity to the nuclear periphery as compared to Chr. 2R-A and Chr. 2R-B (**Figure 1C**). We therefore used the normalized distance between this region and the nuclear periphery as our primary metric for isolating hits.

**Figure 1.**
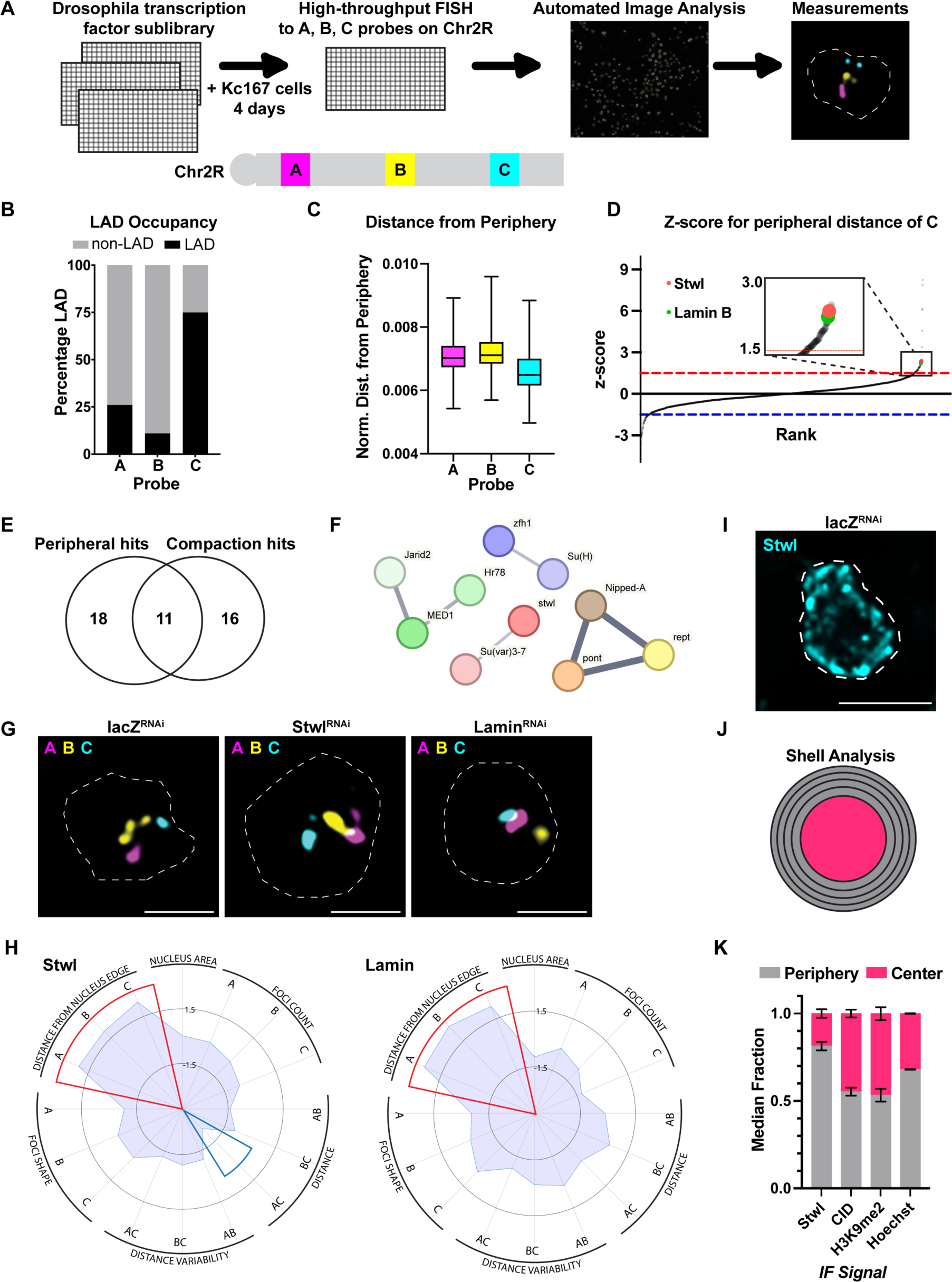
Discovery of novel regulators of chromatin positioning at the nuclear periphery in *Drosophila*. (A) Cartoon schematic of HiDRO screening pipeline and the 1Mb probe regions along chromosome 2R. (B) Percentage of each Chr. 2R region occupied by LADs. (C) Normalized distance from periphery for each Chr. 2R region. (D) Z-score plot for genes affecting peripheral localization of Chr. 2R-C. Genes above red dashed line represent hits that increase the distance between Chr. 2R-C and the nuclear periphery. These are shown larger in the overlay box. Genes below blue dashed line represent hits that decrease the distance between Chr. 2R-C and the periphery. Lamin B and Stwl are shown in green and red, respectively. (E) Venn diagram indicating overlap between the peripheral localization and compaction hits. Eleven genes were hits for both metrics, including stwl. (F) STRING analysis of peripheral hits. (G) Individual Kc167 cell nuclei labelled with probes against Chr. 2R-A (magenta), Chr. 2R-B (yellow) and Chr. 2R-C (blue) from LacZ RNAi (control), Stwl RNAi and Lamin B RNAi. Outlines show nuclear boundary. (H) Radar plot indicating screen metrics following Stwl knockdown (left) or Lamin B knockdown (right). Red and blue wedges represent screen metrics in which the knockdown significantly increased or decreased the metric, respectively. (I) Example nucleus showing Stwl immunofluorescence. Scale bar:5µm. (J) Cartoon schematic of shell analysis of immunofluorescence. Shells 1-4 were combined to define the periphery and shell 5 defines the center. (K) Shell analysis of the indicated nuclear components. The median signal in the periphery and the center was calculated from two replicates of >300 nuclei each.

We performed an RNAi screen in duplicate, using a Drosophila RNAi Screening Center (DRSC)-curated transcription factor dsRNA sublibrary that targets 966 genes encoding DNA-binding or nuclear localizing proteins. A total of ∼8 million cells were analyzed, which yielded 29 “peripheral” hits that significantly increased the distance between Chr. 2R-C and the nuclear periphery, normalized to the nuclear area (**Figure 1D**). In addition to our primary metric, we also calculated peripheral distance for Chr. 2R-A and Chr. 2R-B as well as 13 secondary parameters of genome organization, including the pairwise distance between regions A, B, and C. These also included measurements related to the size and shape of each domain and the nucleus itself, creating a multimodal dataset of nuclear organization for all 966 genes analyzed (**Table S1**). Together, this revealed that 11/29 peripheral hits also altered chromosome length, with all 11 causing increased compaction, consistent with peripheral detachment leading to a global change in genome organization (**Figure 1E**).

### Stwl localizes to the nuclear periphery in Kc167 cells

We used StringDB^36^ to find any known relationships between the peripheral hits and recovered 4 distinct subgroups, one of which included the MADF-BESS domain containing proteins, Su(var)3-7 and Stonewall (Stwl) (**Figure 1F**). Notably, both proteins have been associated with heterochromatin repression^28,33,37,38^. Stwl represented one of our top hits and, similar to lamin B depletion, its phenotypic profile consisted of increased distance for all three Chr2R regions (**Figure 1G-1H**). We also note that Stwl depletion decreased chromosome arm length, as measured by the distance between Chr. 2R-A and Chr. 2R-C.

We next examined the subcellular localization of Stwl in Kc167 cells using an antibody generated against the full-length protein. Reduced immunofluorescence signal from this Stwl antibody following a four-day dsRNA knockdown of Stwl confirmed the specificity of the antibody in Kc167 cells (**Figure S1A-S1C**). Consistent with published reports from other cell types, we found that Stwl was present throughout the nucleus, with an enrichment at the nuclear periphery^33,39^ (**Figure 1I-1K**). Using a shell analysis that divided the nuclear volume into five equi-volume nested shells (**Figure 1I**), we calculated the relative signal in each shell and observed that 82% of Stwl signal occupied the nuclear periphery while 18% occupied the center (**Figure 1J-1K**). In contrast, only 53% of H3K9me2 signal and 55% of CID/CENPA signal occupied the periphery (**Figure 1K**). Total DNA as stained by Hoechst showed only 68% of signal at the periphery (**Figure 1K**), suggesting Stwl was more peripheral than expected for a random distribution throughout the nucleus. We next used affinity purification coupled to quantitative mass spectrometry to determine Stwl interactions in Kc167 cells (**Figure S1D, Table S2**). Consistent with Stwl’s perinuclear localization, we identified putative interactions with multiple components of the nuclear pore complex (NPC) including Nup62 and Nup88. Moreover, we also identified interactions with three other ‘peripheral’ hits from our HiDRO screen, namely Reptin (Rept), Pontin (Pont) and CG4557. Interestingly, Rept and Pont are members of the Ino80 chromatin remodeling complex^40^ and may be required in combination with Stwl to position or repress specific loci at the nuclear periphery. Overall, our data support a direct role for Stwl in anchoring chromatin at the nuclear periphery.

### Stwl promotes perinuclear chromatin positioning independent of Lamin B in Kc167 cells

We next asked if Stwl was required for Lamin expression or localization in Kc167 cells. qPCR and immunofluorescence quantification showed that Lamin B expression was not reduced following Stwl depletion (**Figure S1E-S1F**). To determine if Stwl depletion affected the peripheral localization of Lamin B, we examined 5 distinct Lamin phenotypes and manually assessed >350 cells following LacZ (control) or Stwl depletion (**Figure S1G**). The overall distribution of each phenotype across four Stwl RNAi replicates was not statistically significant from LacZ depletion (**Figure S1H**), suggesting that Stwl relocalizes peripheral chromatin independent of Lamin B expression or localization.

### Stwl regulates chromatin positioning at the nuclear periphery in female GSCs

Stwl has been previously shown to be important for GSC self-renewal, oocyte specification and egg chamber development in *Drosophila* ovaries^28,31,32^. Interestingly, a previous study has also shown that germ cells transform their spatial genome organization during GSC differentiation, including changes in the perinuclear positioning of chromatin^41^. However, the mechanism of Stwl function in GSC maintenance and whether it contributes to GSC genome organization remains unclear. To address this question, we turned to the *Drosophila* ovary, which is a powerful system to study germline stem cell (GSC) fate and tissue homeostasis^42^. Each *Drosophila* ovary comprises 16-20 autonomous egg producing units known as ovarioles. The anterior tip of each ovariole contains a germarium, which houses GSCs and differentiated germ cells (**Figure 2A**). Each GSC divides asymmetrically to produce one self-renewing daughter cell (GSC, green cell) and one differentiating daughter cell (cystoblast, CB, purple cell), with cystoblasts undergoing further transit amplifying divisions to generate germline cysts (yellow cells) (**Figure 2A**). Crucially, the balance between GSC self-renewal and differentiation maintains tissue homeostasis; excessive self-renewal can lead to stem cell tumours while precocious differentiation can lead to tissue atrophy.

**Figure 2.**
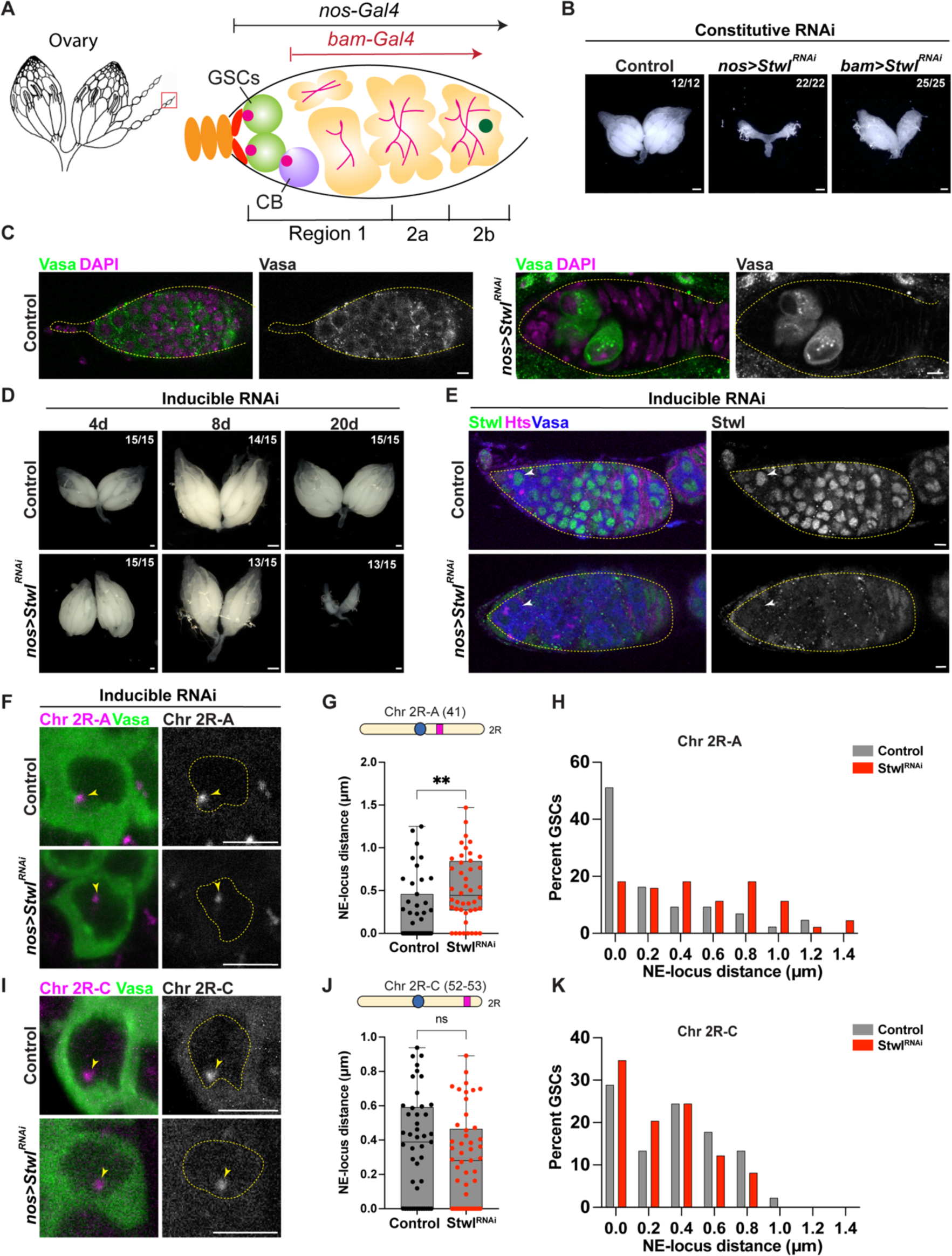
Stwl is a regulator of perinuclear chromatin positioning in female GSCs. (A) Schematic of *Drosophila* ovary and germarium. The germarium resides at the anterior tip of the ovariole (red box) and is further sub-divided into region 1 containing germline stem cells GSCs (green) and cystoblasts, CB (purple) and regions 2a/2b containing differentiated germ cell cysts (yellow). (B) Ovaries from Control *TM3 / Stwl^RNAi^*, nos *> Stwl^RNAi^* and *bam > Stwl^RNAi^* imaged 3 days post eclosion. Scale bar:100μm. (C) Germaria from *nos > mCherry^RNAi^* (Control) and *nos* > *Stwl^RNAi^* ovaries stained for Vasa (green) and DAPI (magenta). Scale bar:5μm. (D) Ovaries from *nos > mCherry^RNAi^* (Control) and *nos* > *Stwl^RNAi^* following 4d, 8d and 20d shift to 29°C in a Gal80^ts^ background. Scale bar:100μm. (E) Germaria from *nos > mCherry^RNAi^* (Control) and *nos* > *Stwl^RNAi^* ovaries stained for Stwl (green), Vasa (blue), and Hts (magenta) following a 4d shift to 29°C. White arrowheads indicate the GSCs. Scale bar:5μm. (F) Oligopaint FISH against Chr. 2R-A (magenta) and IF staining of Vasa (green) in GSCs from *nos > mCherry^RNAi^* (Control) and *nos* > *Stwl^RNAi^* ovaries following a 4d shift to 29°C. Yellow arrowheads indicate the Chr 2R – A locus within the nucleus. Yellow dotted lines indicate the nuclear boundary. Scale bar:5μm. (G) Quantification of NE – Chr. 2R-A distance (µm) in GSCs from (F). n=43 GSCs from *nos > mCherry^RNAi^* and n=44 GSCs from *nos* > *Stwl^RNAi^*. ** indicates p<0.01 from Student’s t-test. (H) Histogram of NE – Chr. 2R-A distance (µm) in GSCs from (G). (I) Oligopaint FISH against Chr. 2R-C (magenta) and IF staining of Vasa (green) in GSCs from *nos > mCherry^RNAi^* (Control) and *nos* > *Stwl^RNAi^* ovaries following a 4d shift to 29°C. Yellow arrowheads indicate the Chr 2R-C locus within the nucleus. Yellow dotted lines indicate the nuclear boundary. Scale bar:5 μm. (J) Quantification of NE – Chr. 2R-C distance (µm) in GSCs from (I). n=45 GSCs from *nos > mCherry^RNAi^*(Control) and n=49 GSCs from *nos* > *Stwl^RNAi^*. ns indicates p>0.05 from Student’s t-test. (K) Histogram of NE – Chr. 2R-C distance (µm) in GSCs from (J).

We first depleted Stwl constitutively in early germ cells (including GSCs and CBs) by RNAi using *nos-Gal4::VP16* (**Figure 2A**). As expected, we observed a severe agametic ovary phenotype upon Stwl depletion (**Figure 2B**) and fully penetrant female sterility (**Figure S2A**). Moreover, these Stwl depleted ovaries contained very few cells containing the germ cell cytoplasmic marker, Vasa (**Figure 2C**). Conversely, although Stwl is expressed in male germ cells, Stwl depletion using *nos-Gal4* in the male germline did not affect testis development or fertility (**Figure S2B-S2D**), suggesting a female germline-specific role for Stwl. Additionally, we verified the Stwl knockdown phenotype using flies carrying a precise *stwl* deletion (*stwl^KO4^*) in trans to a *stwl* mutant allele (*stwl^LL06470^*) (**Figure S2E**). Consistent with the constitutive germline knockdown of Stwl, *stwl* mutant females also exhibited substantial germ cell loss and agametic ovaries (**Figure S2F**). In contrast to the acute loss of early germ cells when Stwl was absent in GSCs, Stwl knockdown in differentiated germ cells using *bam-Gal4* did not affect germaria development (**Figure 2A-2B, Figure S2G-S2H**). Instead, *bam-Gal4*-mediated Stwl depletion led to downstream defects in egg chamber development (**Figure S2I-S2J**), with females exhibiting a strong reduction in fertility compared to controls (**Figure S2K**). The role of Stwl in later stages of oogenesis has been characterized in a separate study^43^. Altogether, these data suggest that Stwl has a critical and cell-autonomous function in female GSC maintenance.

To elucidate the series of events linking Stwl depletion to GSC loss, we used an inducible knockdown system comprising a temperature-sensitive allele of Gal80 (*Gal80^ts^*) and *nos-Gal4*. Here, germ cell specific Gal4 expression is only induced upon shifting the adult flies to 29°C (due to inactivation of *Gal80^ts^*), triggering RNAi and subsequent protein depletion. Using this system, we recapitulated the agametic ovary phenotype observed upon constitutive Stwl depletion 20 days post shift to 29°C (**Figure 2D**). Importantly, we observed that Stwl was depleted in early female germ cells starting from four days post shift to 29°C (**Figure 2E**). Therefore, all further Stwl depletion experiments were performed in flies shifted for 4-6 days to 29°C.

Based on our screen and our phenotypic data in Kc167 cells, we hypothesized that Stwl might position chromatin at the nuclear periphery in female GSCs. Therefore, we first assessed the position of the Chr. 2R regions (A and C) in female GSCs using Oligopaint DNA FISH. Specifically, we measured the shortest distance of these loci from the GSC nuclear boundary, which was marked by the NE-proximal cytoplasmic protein, Vasa. We observed that Chr. 2R-A was positioned closest to the nuclear periphery in control GSCs (median distance = 0 µm, **Figure 2F-2H**). In the absence of Stwl, however, this locus was primarily observed in the nuclear interior (median distance = 0.44µm, **Figure 2F-2G**). In contrast to region A, Chr. 2R-C did not exhibit peripheral localization in control GSCs (median distance = 0.39 µm), consistent with cell-type specific LAD composition, and, as such, its position remained unaffected following Stwl depletion (median distance = 0.28 µm) (**Figure 2I-2K**). We further examined the position of centromeres (marked by the centromeric histone, Cid/ dCENP-A) in GSC nuclei, as they are often observed in proximity to the NE in many cell types^44–46^. While control GSCs exhibited a substantial number of NE-proximal centromeres (**Figure S3A-S3C**), centromeres in Stwl-depleted GSCs were re-localized to the nuclear interior (**Figure S3A-S3C**). Consistent with our data from Kc167 cells, Stwl positions chromatin at the nuclear periphery in female GSCs.

### Reduced peripheral chromatin localization in the absence of Stwl is associated with gaps in the nuclear lamina

We next asked whether Stwl localized to the nuclear periphery in female GSCs as observed in Kc167 cells. We used ovaries enriched for GSC-like cells using *bag-of-marbles* (*bam*) mutants^47^ and stained for Stwl following methanol fixation, a method that can expose otherwise inaccessible epitopes. Interestingly, we observed that a fraction of Stwl consistently localized at the nuclear periphery, interspersed with the nuclear lamina and the nuclear pore complexes (NPCs) (**Figure 3A-3B**), which agrees with our observations in cultured cells. We next sought to identify the underlying cause of the changes in the peripheral chromatin localization observed in Stwl-depleted GSCs. As loss of nuclear envelope integrity is associated with reduced perinuclear chromatin^14–16^, we checked whether Stwl depletion affected NE components in GSCs. We first checked the localization of Lamin B (lamin Dm0) and Lamin C in the Stwl-depleted GSCs since these proteins at the inner nuclear membrane (INM) are associated with peripheral chromatin. We observed that 38% of Stwl-depleted GSCs exhibited stretches of the NE lacking nuclear lamins, referred to as lamina gaps hereafter, with these gaps spanning 10%-40% of the nuclear envelope (**Figure 3C-3D, Figure S4A-S4D**). Importantly, the gaps appeared to be specific to lamins since other INM proteins such as Otefin (*Drosophila* Emerin orthologue) (**Figure S4E-S4F**) and the Lamin B receptor, LBR (**Figure S4G-S4H**) were still present at the gaps. Moreover, we noticed an increased signal intensity of nuclear pore complexes (NPCs) in the lamina gap regions (**Figure 3C-3D**), consistent with previous reports indicating that NPCs can cluster in regions lacking the nuclear lamina^48,49^.

**Figure 3.**
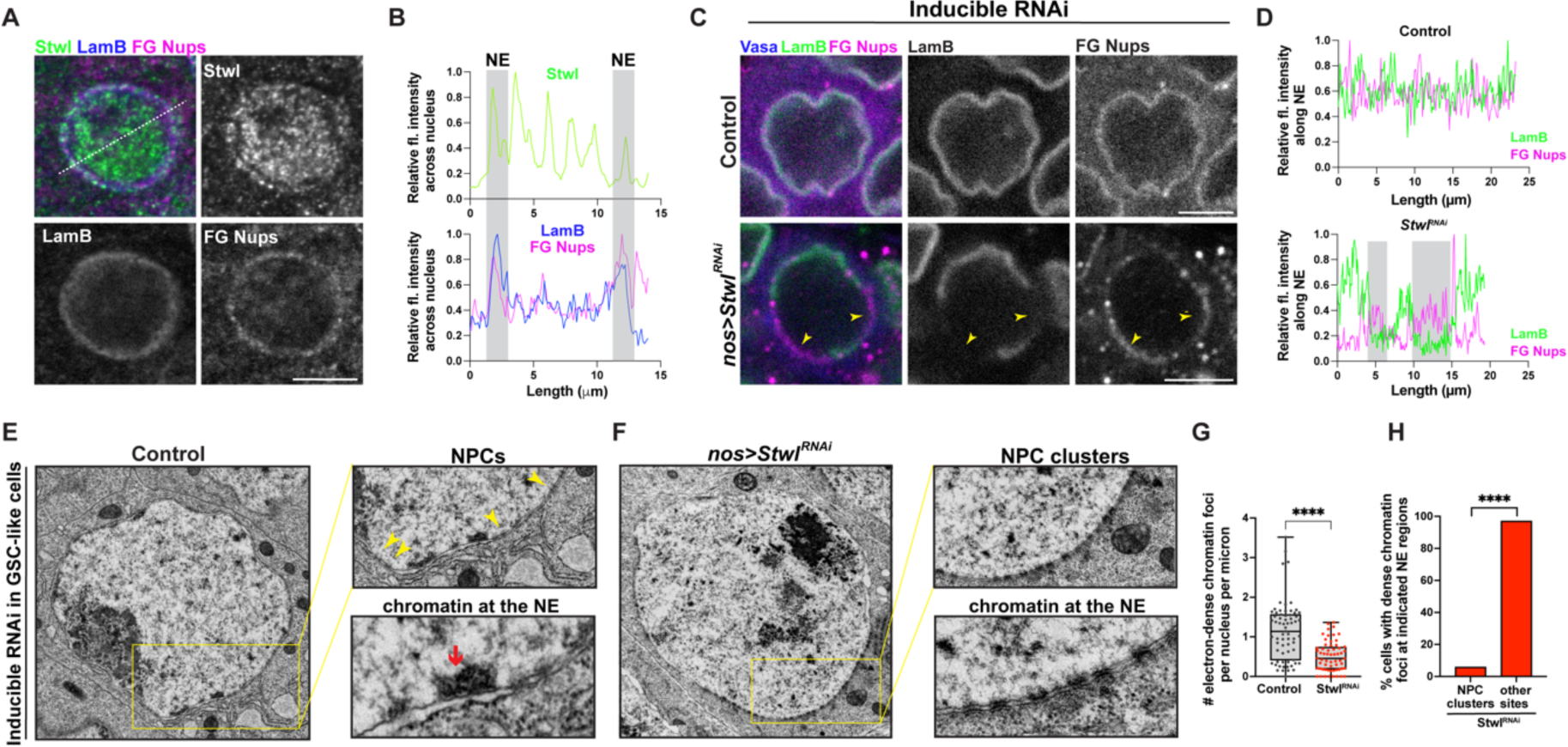
Loss of Stwl leads to defects in perinuclear chromatin organization. (A) IF staining of Stwl (green), Lamin B (blue) and FG Nups (magenta) in GSC-like cells from *bam^Δ86/bam1^* ovaries.Scale bar:5 μm. (B) Relative fluorescence intensity of Stwl (green, top panel), Lamin B (bottom panel, blue) and FG Nups (bottom panel, magenta) across the nucleus (white dotted line) from panel (A). Shaded grey regions highlight the overlap between the three proteins at the NE. (C) IF staining of Lamin B (green), FG Nups (magenta) and Vasa (blue) in GSCs from *nos > mCherry^RNAi^* (Control) and *nos* > *Stwl^RNAi^* ovaries following a 4d shift to 29°C. Yellow arrowheads indicate NPC clusters in the regions lacking Lamin B. Scale bar:5μm. (D) Relative fluorescence intensity of Lamin B (green) and FG Nups (magenta) along the nuclear envelope from (C). Shaded grey regions highlight NPC clustering in regions lacking Lamin B. (E) TEM image of GSC-like cells from *nos > mCherry^RNAi^* (Control) ovaries in a *bam^Δ86^/bam^1^* background following a 4d shift to 29°C. Inset (top) shows NPCs (yellow arrowheads) while inset (bottom) shows an electron-dense chromatin focus associated to the nuclear envelope. (F) TEM image of GSC-like cells from *nos > Stwl^RNAi^* ovaries in a *bam^Δ86^/bam^1^* background following a 4d shift to 29°C. Inset (top) shows NPC clusters while inset (bottom) shows absence of electron-dense chromatin foci in regions containing NPC clusters. (G) Quantification of perinuclear electron-dense chromatin foci in GSC-like cells from (E, F). Each dot represents the number of perinuclear chromatin foci per nucleus per micron of the nuclear envelope. n=67 GSCs from *nos > mCherry^RNAi^* and n=60 GSCs from *nos* > *Stwl^RNAi^*. **** indicates p<0.0001 from Student’s t-test. (H) Percentage of perinuclear electron-dense chromatin foci at NPC clusters versus other regions on the nuclear envelope in GSC-like cells from (F). n=42 **** indicates p<0.0001 from Fisher’s exact test.

To further assess the underlying chromatin ultrastructure at the nuclear periphery, we performed transmission electron microscopy (TEM) in control and Stwl-depleted GSC-enriched ovaries. In contrast to the NE from terminally differentiated mammalian cells, which are lined with compact and electron-dense heterochromatin^1^, *Drosophila* GSCs exhibited multiple distinct perinuclear electron-dense chromatin foci, likely reflecting peripherally localized heterochromatin. In the control, we observed ∼1 electron-dense chromatin focus associated with the nuclear periphery per micron of the nuclear envelope (**Figure 3E-3G**). In contrast, Stwl-depleted GSC nuclei exhibited an approximately 2-fold reduction in the perinuclear electron-dense chromatin foci (**Figure 3F-3G**). In addition, we observed tracts of clustered NPCs in Stwl-depleted GSCs (**Figure 3F**), which likely correspond to the lamina gaps observed by immunofluorescence staining (**Figure 3C-3D**). We next asked whether Stwl depletion led to loss of NPCs from the NE or whether they were rather reorganized across the nucleus. We observed that the normalized number of NPCs (NPCs per micron of the NE) remained unchanged across both control and Stwl-depleted GSCs, suggesting that NPCs are reorganized into clusters in the absence of Stwl (**Figure S4I**). Notably, almost no electron-dense chromatin foci were found in NE stretches with NPC clusters, which correspond to lamina gaps (**Figure 3H**). Consistently, Stwl-depleted GSCs with lamina gaps exhibited fewer NE-proximal centromeric foci in comparison to control GSCs and Stwl-depleted GSCs with an intact lamina (**Figure S3A-S3C**). Taken together, our data suggest that gaps in the nuclear lamina likely contribute to impaired chromatin localization at the nuclear periphery in Stwl-depleted GSCs.

Despite not observing a role for Stwl in Lamin B expression in cultured cells, we considered that reduced levels of Lamin B in Stwl-depleted GSCs could be a possible cause of the lamina gaps and lead to GSC loss. To test this, we used Gal4-mediated *lamin B* overexpression in the female germline. Lamin B overexpression is known to result in cytoplasmic lamin accumulations in *Drosophila* intestinal stem cells (ISCs) and enterocytes (ECs)^50^. Consistently, we observed similar cytoplasmic lamin accumulations following *lamin B* over-expression in GSCs (**Figure S5A-S5B**). However, Lamin B overexpression in Stwl-mutant GSCs failed to rescue the lamina gaps or the atrophied ovary phenotype (**Figure S5C-S5D**). These data suggest that a decrease in Lamin B protein levels is not a primary cause of lamina gaps and GSC loss in the absence of Stwl.

Recent reports have shown that loss of the INM protein, Otefin, triggers a *Chk2*-dependent GSC developmental arrest in *Drosophila* ovaries, with *Chk2* mutation partially restoring germline development in the absence of Otefin^51^. However, *Chk2* and *Stwl* double mutants did not rescue GSC loss or ovary atrophy (**Figure S5E**). Finally, we tested whether germ cell death markers such as lysotracker and Death caspase 1 (Dcp-1) were elevated in Stwl-depleted germaria^52^. While we did observe cell death in the absence of Stwl, the death was restricted to differentiated germline cysts and not observed in GSCs (**Figure S5F-S5G**). Thus, our data point to a distinct mechanism for GSC loss and ovary atrophy in the absence of Stwl.

### Stwl represses the expression of the GSC differentiation gene, *benign gonial cell neoplasm* (*bgcn*)

Based on our FISH and TEM data, we hypothesized that loss of peripheral chromatin organization in the absence of Stwl might contribute to GSC loss through altered transcriptional programs. To test this, we first wanted to identify the Stwl-dependent transcriptome, specifically in GSC-like cells. Although other studies have identified Stwl-dependent gene expression in ovaries, these studies were performed in young ovaries that contain early egg chambers, differentiated germline cysts as well as GSCs^34,43^. Moreover, Stwl-depleted ovaries rapidly lose GSCs (**Figure 2B-2C**) and are therefore unsuitable for RNA-seq experiments that seek to determine the GSC transcriptome. However, a previous study has shown that overexpression of Stwl in GSCs leads to a subtle increase in the number of undifferentiated germ cells in the ovary^31^. Interestingly, we found that Stwl overexpression further enhanced the number of undifferentiated (Bam-negative) germ cells in a *bam* heterozygous background, where GSC differentiation signaling is likely weakened (**Figure 4A-4B**). This data further strengthens the idea that Stwl overexpression can promote GSC fate. We therefore performed RNA-seq to identify Stwl-dependent genes in control and Stwl overexpressing (Stwl^OE^) GSC-enriched ovaries. We observed 548 genes differentially expressed following Stwl overexpression (log_2_FC>|0.6|, p_adj_<0.01), with 154 genes downregulated in comparison to the control (**Figure 4C, Table S3, Table S4**). We specifically focused on the downregulated genes since Stwl is reported to function as a transcriptional repressor^28,31,33^. Here, we found that the expression of the GSC differentiation gene, *benign gonial cell neoplasm* (*bgcn*)^53–55^, loss of which results in the accumulation of undifferentiated GSC-like cells in the *Drosophila* ovary, was reduced 1.9-fold upon Stwl overexpression (**Figure 4C-4D**). In addition, we identified that an inhibitor of ecdysone signaling, the transcriptional corepressor, *SMRT-related and ecdysone receptor interacting factor* (*Smr*)^56^, was also downregulated 1.5-fold following Stwl overexpression (**Figure 4C-4D**). Since ecdysone signaling is critical for GSC self-renewal and maintenance^57^, Smr activity may promote differentiation and is likely repressed in GSCs. Interestingly, both *bgcn* and *Smr* are upregulated in gene expression datasets from other studies using Stwl-depleted ovaries^31,34,43^, suggesting that these genes are likely repressed in a Stwl-dependent manner (**Figure 4C**). Furthermore, we performed Cleavage Under Targets and Release Using Nuclease (CUT&RUN)^58^ chromatin profiling experiment in GSC-enriched ovaries to identify the direct targets of Stwl (**Figure 4E**). We observed Stwl peaks mostly at non-coding sequences such as promoters (∼47%) but also at introns, UTRs and distal intergenic regions (**Figure 4E and Figure S6A**). We next assessed the extent of overlap between Stwl bound loci in GSCs and differentially expressed genes upon Stwl overexpression. We found that 59.1% of downregulated genes and 69.5% of upregulated genes had a Stwl peak within 1kb of the gene body (**Figure S6B**). Importantly, Stwl was bound to genomic regions in close proximity to the *bgcn* and *Smr* gene loci (**Figure 4F and Figure S6C**), further indicating that Stwl may directly bind and regulate the expression of these genes.

**Figure 4.**
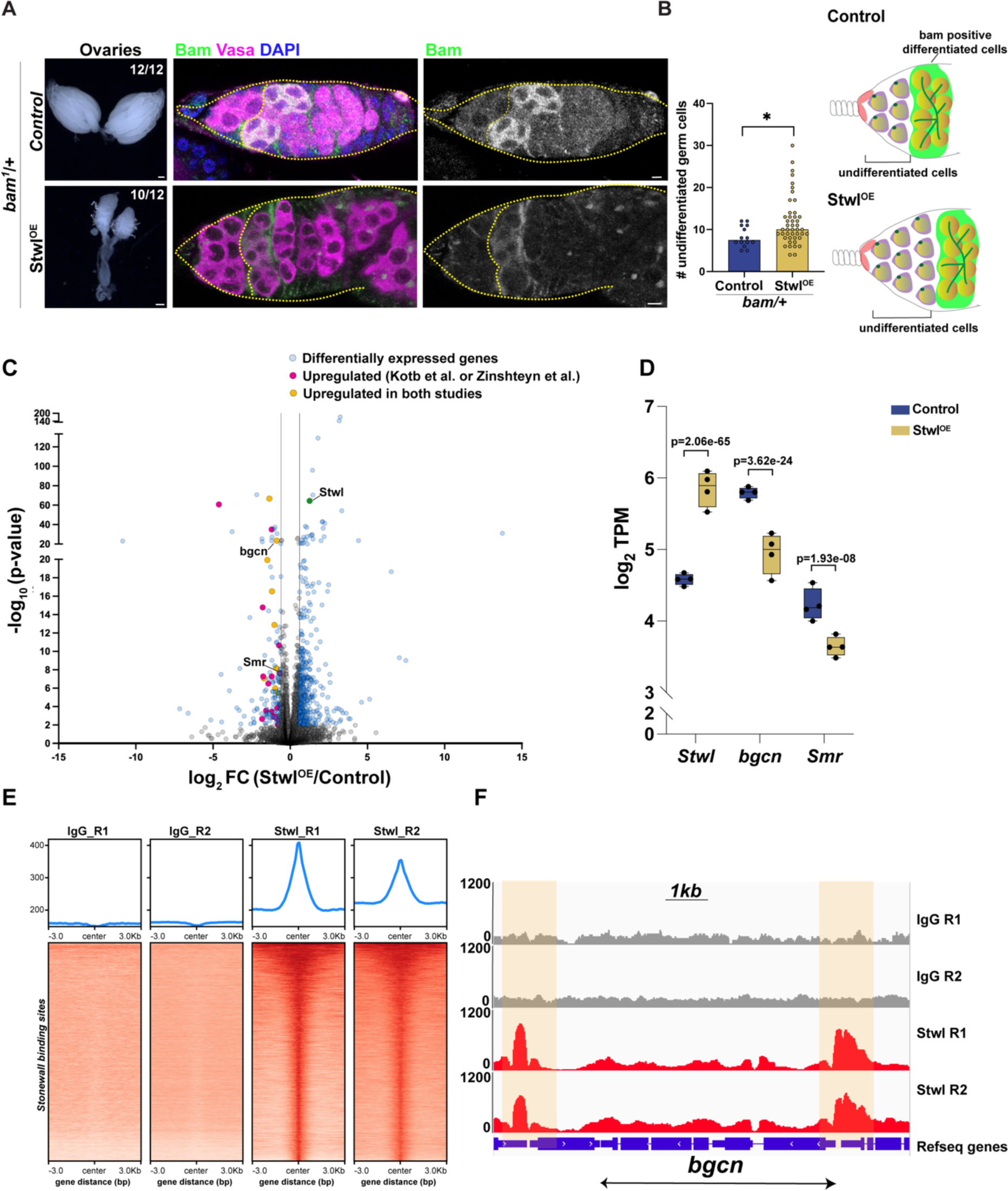
Stwl binds and represses the *bgcn* differentiation gene. (A) First panels, *nos*; *TM3* (Control) and *nos* > *Stwl^EY00146^* (*Stwl^OE^*) ovaries in a *bam*^1^/*+* background. Middle and right panels, IF staining of Bam (green), Vasa (magenta) and DAPI (blue) in germaria. Scale bar:5μm (B) Quantification of undifferentiated Bam-negative germ cells from (A). n=15 germaria from the control and n=45 germaria from *Stwl^OE^*. * indicates p<0.05 from a Student’s t-test. (C) Volcano plot of −log_10_(p-value) vs log_2_FC from *nos-Gal4*/+ (Control) and *nos* > *Stwl^EY00146^* (*Stwl^OE^*) GSC-enriched ovaries (*bam^Δ86^/bam^1^* background). Differentially expressed genes (log_2_FC>|0.6| and p_adj_<0.01 are indicated as blue dots. Genes upregulated in Stwl-depleted ovaries from Zinshteyn et al.^34^ or Kotb et al., 2023^43^ are indicated as magenta dots while genes upregulated in both studies are indicated as yellow dots. Adjusted p values following multiple testing correction are shown. (D) Transcripts per million (log_2_TPM) for the indicated genes from *nos-gal4*/+ (Control) and *nos* > *Stwl^EY00146^* (*Stwl^OE^*) GSC-enriched ovaries in a *bam^Δ86^/bam^1^* background. Adjusted p values following multiple testing correction are shown. (E) Heatmaps of CUT&RUN reads for IgG from young WT ovaries and for Stwl from ovaries enriched for GSC-like cells (*nos* > *bam^RNAi^*). Data are centered on ±3 kb window around 12888 Stwl peaks (merged within 1kb) and is shown for two replicates each. (F) Capture of the IGV genome browser (v2.11.4) showing an approximately 10kb region on *Drosophila* chromosome 3 (y axis = reads per kilobase per million reads). Ensembl genes (blue). Shaded areas correspond to Stwl binding peaks.

### Stwl positions *bgcn* at the nuclear periphery to regulate its expression

Our data thus far indicate that Stwl can position chromatin at the nuclear periphery in GSCs and repress GSC differentiation genes such as *bgcn*. To test whether these two functions of Stwl were linked, we first assessed the position of the *bgcn* locus in relation to the GSC nuclear periphery using Oligopaint DNA FISH. We measured the shortest distance between the *bgcn* locus and the nuclear periphery in control and Stwl-depleted GSCs. We observed a 1.7 fold reduction in *bgcn* loci at the nuclear periphery of GSCs in the absence of Stwl (control GSCs – 34% peripheral *bgcn* loci, Stwl-depleted GSCs – 20% peripheral *bgcn* loci, **Figure 5A-5C**). We also assessed the position of the *bgcn* locus in differentiated germline cysts within the germarium (Region 2a/2b, **Figure 2A**). Similar to GSCs, we observed a 1.5 fold reduction in peripherally localized *bgcn* loci in Stwl-depleted germline cysts (control cysts – 61% peripheral *bgcn* loci, Stwl-depleted cysts – 39% peripheral bgcn loci, **Figure 5D-5F**). The increased peripheral localization of *bgcn* loci in differentiated germline cysts in comparison to GSCs (61% in germline cysts vs 34% in GSCs) is consistent with the observation that *bgcn* expression is typically only observed in GSCs and CBs^53^. Importantly, Stwl promotes *bgcn* positioning at the nuclear periphery in both GSCs and differentiated germline cysts.

**Figure 5.**
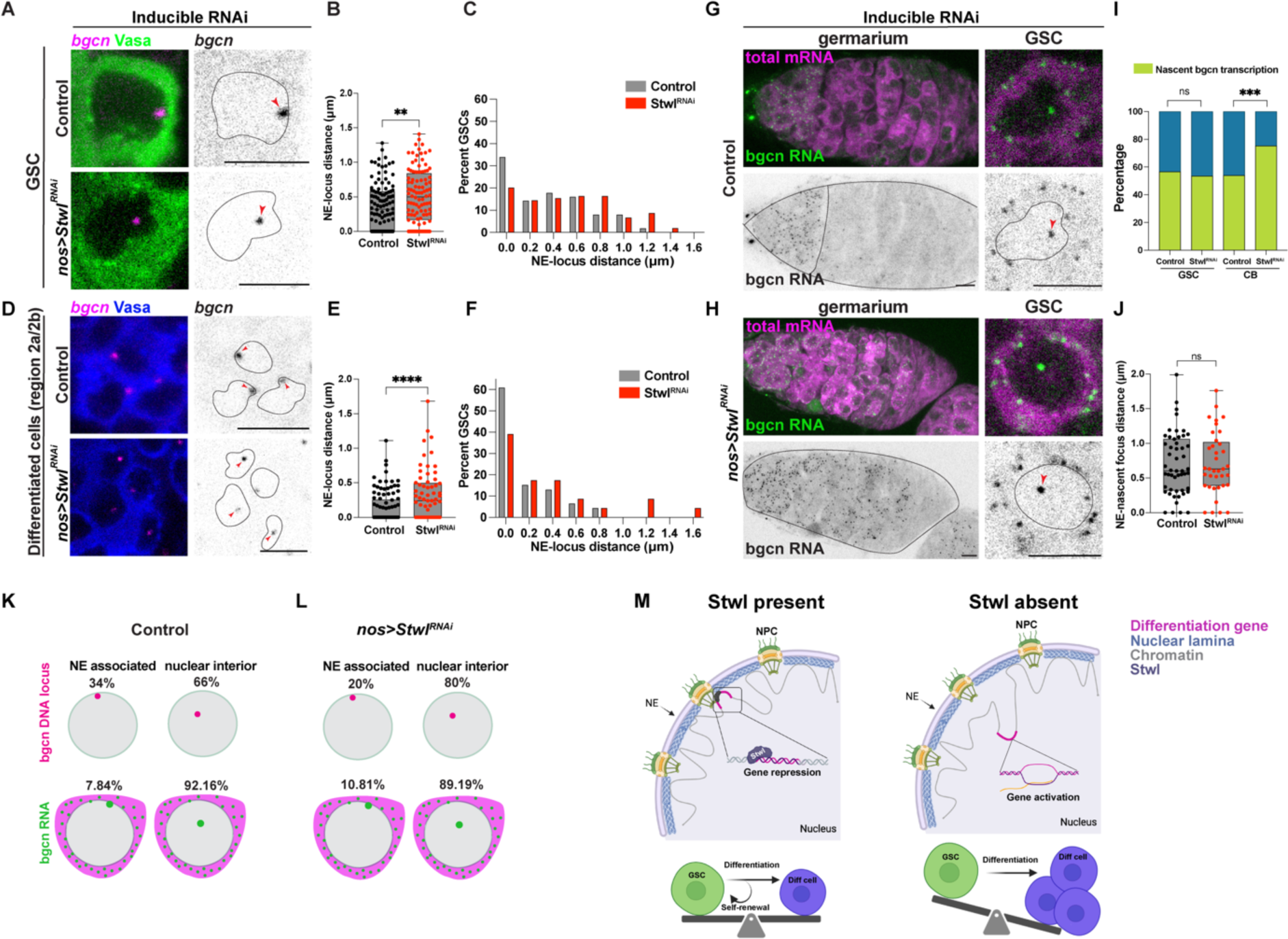
Stwl positions *bgcn* at the nuclear periphery in female GSCs. (A) Oligopaint FISH against the *bgcn* locus (magenta) and IF staining of Vasa (green) in GSCs from *nos > mCherry^RNAi^*(Control) and *nos* > *Stwl^RNAi^* ovaries following a 6d shift to 29°C. Red arrowheads indicate the *bgcn* locus within the nucleus. Black dotted lines indicate the nuclear boundary. Scale bar:5 μm. (B) Quantification of NE – *bgcn* distance (µm) in GSCs from (A). n=112 GSCs from *nos > mCherry^RNAi^* (Control) and n=104 GSCs from *nos* > *Stwl^RNAi^*. ** indicates p<0.01 from Student’s t-test. (C) Histogram of NE – *bgcn* distance (µm) in GSCs from (B). (D) Oligopaint FISH against the *bgcn* locus (magenta) and IF staining of Vasa (blue) in region 2a/2b differentiated germline cysts from *nos > mCherry^RNAi^* (Control) and *nos* > *Stwl^RNAi^* ovaries following a 6d shift to 29°C. Red arrowheads indicate the *bgcn* locus within the nucleus. Black dotted lines indicate the nuclear boundary. Scale bar:5μm. (E) Quantification of NE – *bgcn* distance (µm) in region 2a/2b differentiated germ cells from (D). n=106 GSCs from *nos > mCherry^RNAi^* (Control) and n=65 GSCs from *nos* > *Stwl^RNAi^*\*\*\*\* indicates p<0.0001 from Student’s t-test. (F) Histogram of of NE – *bgcn* distance (µm) in region 2a/2b differentiated germ cells from (E). (G) smFISH against *bgcn* mRNA (green) and poly-A mRNA (magenta) in GSCs from *nos > mCherry^RNAi^* (Control) following a 6d shift to 29°C. In left panel, black dotted lines demarcate region 1 and the germarium boundary. In right panel, black dotted lines indicate the nuclear boundary. Scale bar:5 μm. (H) smFISH against *bgcn* mRNA (green) and poly-A mRNA (magenta) in GSCs from *nos > Stwl^RNAi^* following a 6d shift to 29°C. In left panel, black dotted lines demarcate the germarium boundary. In right panel, black dotted lines indicate the nuclear boundary. Scale bar:5 μm. (I) Quantification of percentage of GSCs and cystoblasts (CBs) with nascent bgcn expression from *nos > mCherry^RNAi^* (Control) and *nos* > *Stwl^RNAi^* ovaries following a 6d shift to 29°C. For the control, n=90 (GSCs) and n=174 (CBs). For *nos* > *Stwl^RNAi^*, n=71 (GSCs) and n=146 (CBs). ns indicates p>0.05 and *** indicates p<0.001 from a Fisher’s exact test. (J) Quantification of NE – *bgcn* nascent focus distance (µm) in GSCs from (G, H). n=51 GSCs from *nos > mCherry^RNAi^* (Control) and n=37 GSCs from *nos* > *Stwl^RNAi^*. ns indicates p>0.05 from Student’s t-test. (K, L) Schematic of data from (A – C and G – H) showing the percentage of GSCs with the *bgcn* DNA locus and the nascent *bgcn* RNA focus positioned at the nuclear periphery or in the nuclear interior in GSCs from *nos > mCherry^RNAi^* (Control) and *nos* > *Stwl^RNAi^* ovaries following a 6d shift to 29°C. (M) Model for Stwl function in female germline stem cells.

Does the position of the *bgcn* gene within the nucleus dictate its expression? To address this question, we performed single molecule RNA FISH (smFISH) in control and Stwl-depleted ovaries. We used FISH probes targeting *bgcn* exons, which mark cytoplasmic *bgcn* mRNA molecules as well as nascent transcripts emanating from the *bgcn* gene locus. In control cells, cytoplasmic *bgcn* transcripts were primarily observed in the GSCs and cystoblasts (**Figure 5G**), consistent with previous reports^53^. In contrast, Stwl-depletion resulted in cytoplasmic *bgcn* transcripts across the entire germarium, including differentiated germline cysts in region 2a/2b (**Figure 5H**). Importantly, we observed significantly more CBs and differentiated germline cysts with nascent *bgcn* transcription upon Stwl depletion (**Figure 5I**), which strongly correlates with reduced frequency of *bgcn* loci at the nuclear periphery (**Figure 5A-5F**).

Strikingly, nearly all *bgcn* nascent transcription in control and Stwl-depleted GSCs was observed in the nuclear interior (**Figure 5J**). For example, although 34% of *bgcn* gene loci are perinuclear in control GSCs, *bgcn* nascent transcription was predominantly observed in the nuclear interior (92% of control GSCs with *bgcn* nascent transcription, **Figure 5K**). This suggests that the majority of perinuclear *bgcn* gene loci are transcriptionally silent. We observed a similar effect in Stwl-depleted GSCs, where *bgcn* nascent transcription was again primarily observed in the nuclear interior (89% of Stwl-depleted GSCs with *bgcn* nascent transcription, **Figure 5L**). Thus, the *bgcn* loci that remain at the nuclear periphery are not transcribed, even in Stwl-depleted GSCs. These data suggest that the primary function of Stwl may be to position specific chromatin loci or genes at the nuclear periphery, where they are kept transcriptionally silent through the action of other factors. Taken together, we propose a model where Stwl promotes GSC fate through perinuclear positioning and repression of differentiation genes such as *bgcn*.

## Discussion

The regulation of gene expression is a primary mechanism that dictates cell fate. In addition to local factors influencing gene expression such as enhancer-promoter contacts and sequence-specific transcription factors, the position of a gene within the nucleus can also influence expression^1,13,59^. In many organisms, the enrichment of dense and compact heterochromatin at the nuclear periphery gives rise to a gene-repressive nuclear subcompartment. Consistently, genes anchored to the nuclear periphery are generally transcriptionally inactive while repositioning the same genes to the nuclear interior is associated with their expression^1,13^. In many species, INM-associated proteins and repressive chromatin modifications mediate large-scale chromatin tethering to the nuclear envelope^1,2,13^. However, chromatin-associated proteins that position specific gene loci at the nuclear periphery are largely unidentified, even in powerful multicellular model organisms such as *Drosophila*.

In this study, we have deployed HiDRO^27^ in tandem with a high-throughput RNAi screen for factors influencing nuclear architecture in *Drosophila*. We have identified 29 hits affecting chromatin positioning at the nuclear periphery, including multiple heterochromatin-associated proteins such as Su(var)3-7, HP2 and Jarid2 as well as transcription factors such as Su(H), Sry-delta and Fer2, with many of these hits known to have important roles in specific cell types^38,60–63^. Among these hits, we have revealed that Stonewall (Stwl), a MADF-BESS transcriptional regulator previously implicated in female GSC maintenance^28,31,32^, is a novel factor positioning chromatin at the nuclear periphery in *Drosophila* cultured cells and female GSCs (**Figure 5M**). Using a multimodal approach, we identify that Stwl binds and represses many genes in female GSCs, including canonical differentiation genes such as *bgcn* as well as genes implicated in differentiation such as *Smr*. We propose that Stwl-mediated repression of multiple such genes through perinuclear positioning preserves the balance between self-renewal and differentiation, thereby ensuring the long-term maintenance of the GSC reservoir and preserving tissue homeostasis (**Figure 5M**).

Although the nuclear periphery is considered to be a repressive nuclear subcompartment^1^, whether perinuclear gene position dictates transcriptional activity or whether transcriptional activity drives perinuclear positioning of genes has remained incompletely understood. Our identification of novel perinuclear anchors such as Stwl, and the genomic loci that they bind and repress, highlights a path forward to address this challenging question. For example, oligopaint DNA FISH experiments revealed that the Stwl-bound *bgcn* gene locus was often positioned at the nuclear periphery in GSCs and differentiated germline cysts. This perinuclear positioning was reduced 1.5-1.7-fold in the absence of Stwl and was broadly associated with increased *bgcn* expression across the germaria as detected by smFISH. Interestingly, *bgcn* nascent transcription was primarily observed in the nuclear interior and rarely observed at the nuclear periphery in the same cell types. We observed a similar lack of *bgcn* nascent transcription at the nuclear periphery in GSCs lacking Stwl. Since the *bgcn* locus is present at the nuclear periphery in 34% and 20% of control and Stwl-depleted GSCs respectively, our data are consistent with a model where Stwl primarily functions to position loci at the nuclear periphery, and that other components of the perinuclear heterochromatin subcompartment mediate direct transcriptional repression. However, Stwl may also have other complementary roles that facilitate transcriptional repression at bound loci. For example, Stwl may possess a direct transcriptional repression activity that only operates at the nuclear periphery, potentially through interactions with specific NE-associated proteins.

Cytologically, we observed that a fraction of Stwl localizes to the nuclear periphery in both cultured *Drosophila* cells and female GSCs. Moreover, we identified interactions between Stwl and NPC proteins (Nup62, Nup88, Nup214 and Tpr/Megator) through quantitative proteomics in cultured cells. While it is possible that these interactions could facilitate nuclear import of Stwl, recent studies have also shown that the perinuclear localization of active and repressive chromatin can occur through interactions with NPC proteins^19–21^. Interestingly, we also identified that three other ‘peripheral’ hits from our screen (Reptin, Pontin and CG4557) co-purified with Stwl, suggesting that a Stwl-containing multi-protein complex may be required to facilitate perinuclear positioning of bound loci. At the same time, our discovery of multiple potential perinuclear anchors suggests a high degree of redundancy in the system. One example of this potential redundancy may be the male germline, where Stwl depletion has no effect on GSC maintenance. We therefore speculate that other proteins function in parallel to Stwl outside of the female germline, and these proteins may include other ‘peripheral’ hits identified in our screen (e.g. Jarid2 and Su(var)3-7) or one of the 45 *D. melanogaster* MADF-BESS family members (e.g. Brwl or Hng2)^29,64^, which are known to function redundantly in other tissues.

In the absence of Stwl, we observe that GSCs undergo substantial changes in chromatin organization at the nuclear envelope, including decreased electron-dense perinuclear chromatin foci and gaps in the nuclear lamina. The decreased perinuclear chromatin association in the absence of Stwl could be due to a lack of bridging interactions between chromatin and the nuclear envelope. However, another possibility is that Stwl-dependent genome organization may also promote perinuclear chromatin association. A parallel study^43^ has identified that Stwl is enriched at the boundaries between active and inactive genomic regions in young ovaries, in a manner reminiscent of insulator proteins that demarcate topologically associated domains (TADs)^65^. In the absence of Stwl, they find that the chromatin states of these active-inactive regions are indistinct, which is suggestive of compartment mixing and is associated with gene misexpression. Intriguingly, previous studies have noted that transcriptionally silent lamina-associated domains (LADs) are separated from neighbouring active genomic compartments by a sharp border^4,66^. In addition, induced expression of peripherally positioned genes and alteration of their chromatin state results in relocalization to the nuclear interior^67,68^. Kotb and colleagues have further shown that mixing of active and inactive chromatin states at the *Rps19b* locus in the absence of Stwl is associated with detachment from the nuclear periphery in nurse cells^43^. Therefore, we postulate that heterochromatin-euchromatin compartment mixing in the absence of Stwl may destabilize heterochromatin domains and perinuclear chromatin anchoring.

In summary, our HiDRO-based nuclear architecture screen has identified multiple potential chromatin-associated perinuclear anchors in the *Drosophila* genome. Here, we have focused on Stwl, which we identified as a factor required for positioning chromatin at the nuclear periphery in female GSCs. Strikingly, we show that this property of Stwl is critical to promote female GSC fate, through the anchoring of canonical differentiation genes at the repressive perinuclear sub-compartment. Thus, our study makes a significant step toward dissecting causal relationships between the position of a gene, the regulation of its expression and the effect on cell fate decisions in multiple tissues.

## Materials and Methods

### *Drosophila* husbandry and strains

All flies were raised on standard Bloomington medium at 25°C unless otherwise noted. *Stwl^RNAi^* (BDSC35415), *mCherry^RNAi^* (BDSC35785), *P{EPgy2}stwl^EY00146^* (BDSC21350), *bam^Δ86^*(BDSC5427), *bam^RNAi^* (BDSC33631) were obtained from the Bloomington Drosophila Stock Center. *stwl^LL6470^* (DGRC141809) was obtained from the Kyoto Stock center. *nos-GAL4 ^+VP16^* (3^rd^ chromosome)^69^*, bam-GAL4*^70^ and *bam*^1^ ^71^ have been previously described. *nos-GAL4 ^+VP16^* (2^nd^ chromosome) and *nos-GAL4 ^−VP16^*; *Gal80^ts^* were gifts from Yukiko Yamashita. For inducible knockdown experiments, *nos-GAL4 ^−VP16^*; *Gal80^ts^* flies were crossed to the desired RNAi strain at 18°C. Following eclosion, 1 day old flies were collected and shifted to 29°C to induce RNAi expression.

### HiDRO and screen data analysis

HiDRO was adapted from Park et al., 2023^27^ for Drosophila cells. 384-well plates (Perkin Elmer #6057300) were seeded with dsRNA by the DRSC screening core at Harvard. Kc167 cells were resuspended at a concentration of 1×10^6^ cells/ml in serum-free Schneider’s S2 media (Thermo Fisher #R69007) and seeded onto 384-well plates at a volume of 10 µl per well using the Matrix WellMate (Thermo Fisher) and then spun down at 1200 rpm for 2 min. Unless otherwise indicated, spins were done at this setting and pipetting was performed by the WellMate. Plates were allowed to incubate at 25°C for 30 minutes for dsRNA uptake before being seeded with 30 µl of serum-containing media. Cells were allowed to grow for 4 days. To fix the cells, cells were first washed with 1x PBS and then fixed in 4% paraformaldehyde in 1x PBS for 5 min, with plates spun right after the addition of the fixative to ensure full contact with the cells. PFA was removed and cells were washed and stored in 1x PBS at 4°C.

For the first day of the FISH protocol, 1x PBS was used to wash the cells prior to the addition of a solution of 50% formamide in 2xSSC and 0.1% Tween-20 (50%FMM/2xSSCT). Plates were spun and then incubated at 91°C for 3 min on heat blocks (VWR), then 60°C for 20 min, and then allowed to cool to room temperature. Wells were aspirated and then filled manually with a multichannel pipette with 20 µl of hybridization mix containing 50%FMM/2xSSCT and 1 pmol of each probe. Plates were spun and placed on the heat blocks for 20 min at 91^°^C. Plates were spun one more time before incubating on the hot block overnight at 37°C.

For the second day, plates were washed several times with 2xSSCT to completely remove the hybridization mix from wells. Then, plates were incubated twice with 2xSSCT prewarmed at 60°C for 5 min. Plates were then incubated with room temperature 2xSSCT for 5 min, with the last wash containing 1 µg/ml of Hoechst 33342. Then, plates were incubated twice with room temperature 2xSSC for 15 min prior to the addition of imaging buffer containing 2x SSC, 10% glucose, 10 mM Tris-HCl, 0.1 mg/mL catalase, 0.37 mg/mL glucose oxidase.

Plates were imaged within 5 days of the FISH protocol on the Yokogawa CV7000 at the NCI High-Throughput Imaging Facility (HiTIF) with the 60x objective, and 2×2 pixel binning to achieve a resolution of 0.217 µm per pixel. 10 fields were imaged per condition, with Z-stacks consisting of 21 slices at 0.5 µm intervals imaged and max-projected for 2D analysis.

Images from HiDRO plates were segmented and measured using CellProfiler v3.1.8^72^.Both nuclei and FISH foci were identified using the “global” thresholding strategy and the “Otsu” method. All metrics from the “MeasureObjectSizeShape” module were exported and processed as follows. First, measurements from individual nuclei were summarized by determining the minimum distance of spots to the nuclear periphery, the minimum distance between spots, and the average eccentricity value for each spot. Then, data from all the nuclei per well were aggregated by averaging, and z-scores were calculated by comparing the well average to the distribution of values of all wells of the same plate. In order for a gene to be considered a hit, at least two replicates of the same dsRNA treatment for that gene had to surpass an absolute z-score cutoff equal to or larger than of 1.5.

### dsRNA production

The following primers were used to both amplify the gene of interest from genomic DNA and add T7 adapters. The resulting PCR products were purified using a NucleoSpin Gel and PCR Cleanup kit (Macherey-Nagel). dsRNA was generated using the MEGAscript T7 kit (Invitrogen) and purified using the RNeasy kit (Qiagen). dsRNAs were heated to 65°C for 30 minutes and then cooled slowly to room temperature to renature dsRNA.

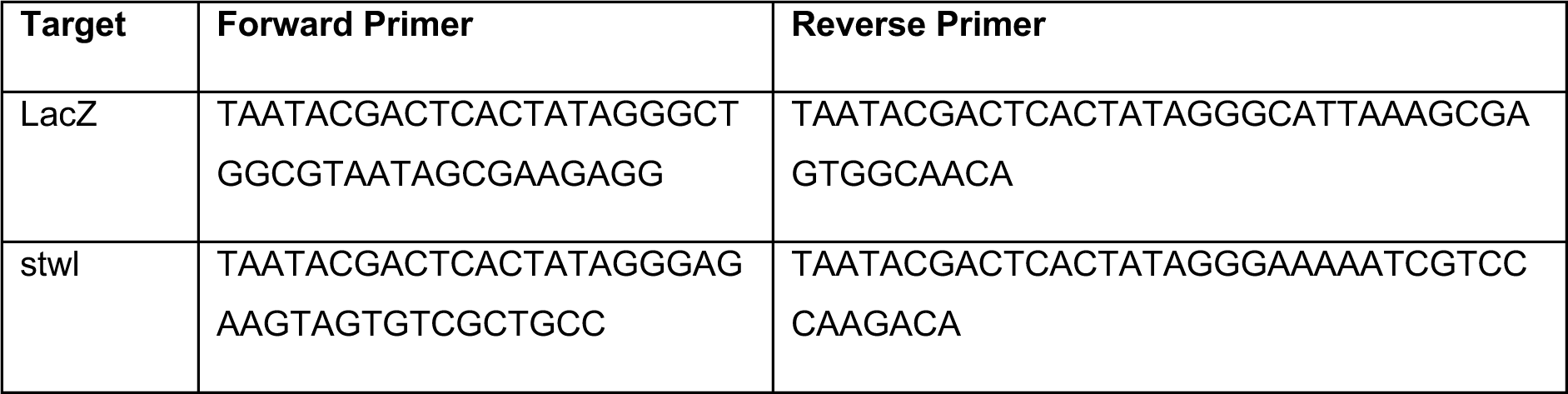

### Cell culture and knockdowns

Kc167 cells were obtained from the *Drosophila* Genome Resource Center (DGRC). Cells were grown at 25°C in Schneider’s medium, supplemented with 10% FBS. Cultures were split twice per week at a 1:4 ratio. For knockdowns, 4×10^6^ cells were incubated with 40µg of dsRNA in 1mL of serum-free medium for 30 mins in each well of a six-well plate. After incubation, 3mL of complete medium was added to the cells. Cells were cultured for four days. Control cells were treated with dsRNA targeting LacZ.

### qPCR

RNA was extracted from cells using the RNeasy Kit (Qiagen) and converted to cDNA using the Maxima Reverse Transcriptase kit (Thermo Scientific). qPCR was run using PowerUp SYBR Green Master Mix (Applied Biosystems). Genes of interest were compared to the geometric mean of three housekeeping genes (Aldh7A1, P5CS, and Ssadh). Primers used are listed in the table below.

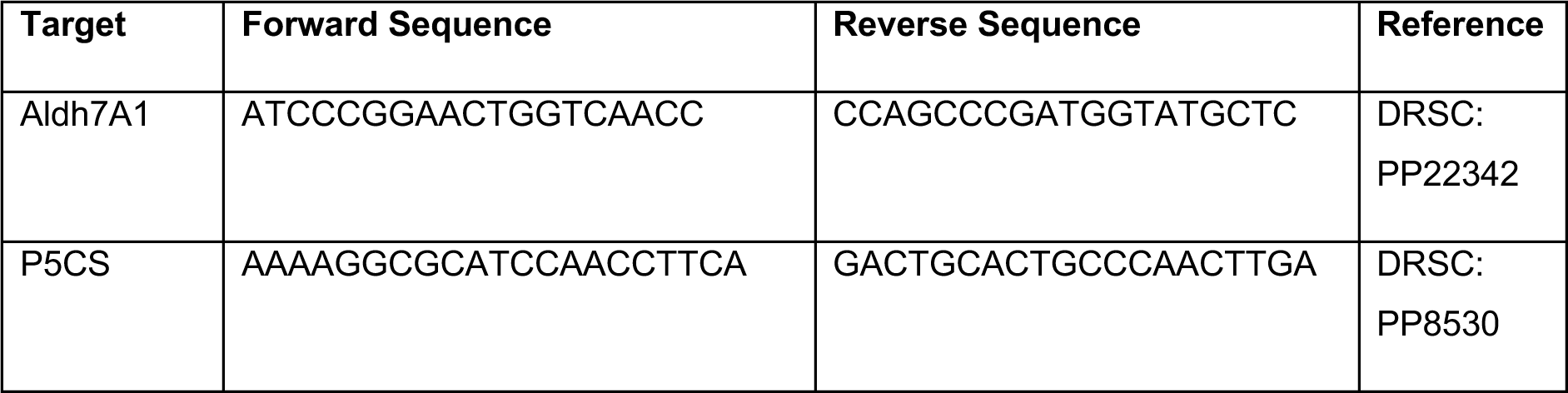
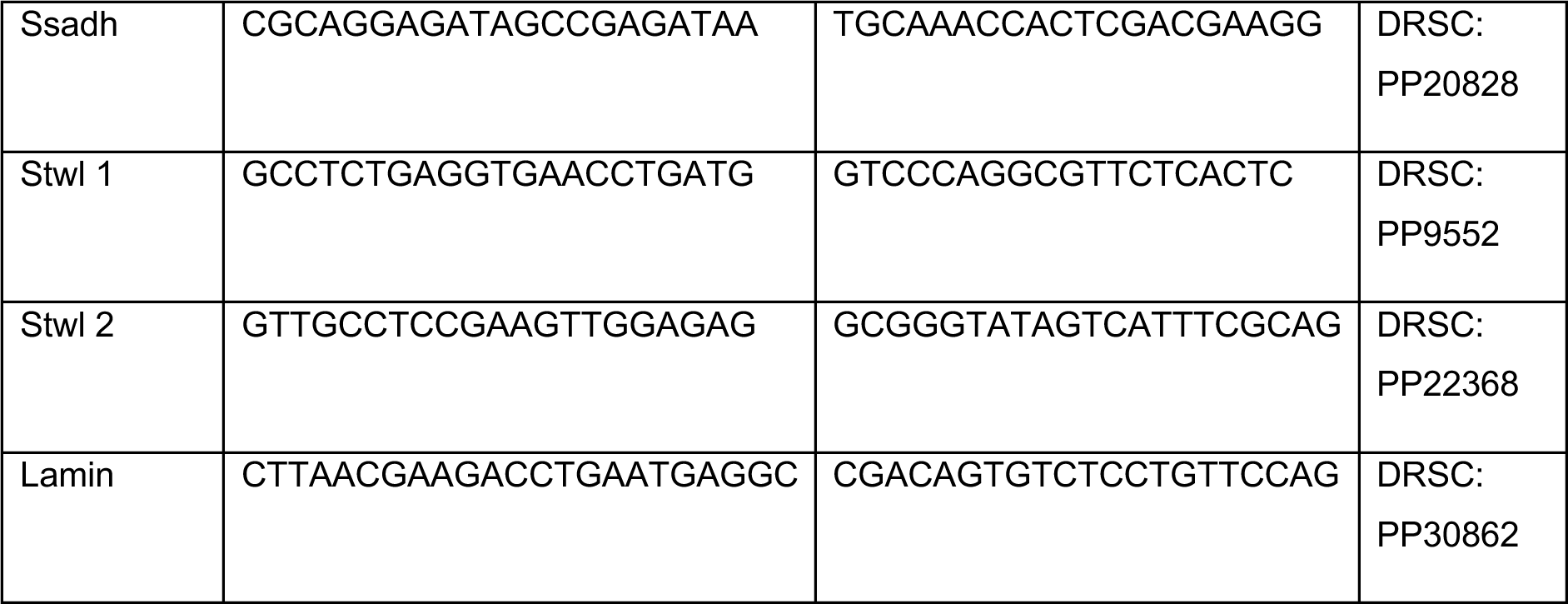

### Purification of Stonewall for antibody generation

For expression of N-terminal His_6_-tagged Stwl in bacteria, the stonewall coding sequence was amplified by PCR and cloned into the XhoI and NcoI sites of the pET28a vector (Novagen). The plasmid was transformed into *E.coli* BL21(DE3) cells (StrataGene) and protein expression induced with 0.5 mM IPTG at 37°C for 4 h. For protein purification, cells were resuspended in lysis buffer (6 M GndHCl, 0.1 M NaH_2_PO_4_ and 0.01 M Tris-HCl (pH 8.0)), followed by incubation at RT for 60 min. The lysate was cleared by centrifugation at 12’000 g for 30 min at RT and added to Ni-NTA agarose beads (Qiagen) equilibrated in lysis buffer. After incubation for 1 h at RT, beads were washed once with lysis buffer and twice with wash buffer (8 M Urea, 0.1 M NaH_2_PO_4_ and 0.01 M Tris-HCl (pH 6.3)). His_6_-Stwl was eluted with wash buffer adjusted to pH 4.5 and rebuffered to 1X PBS by dialysis. Antibodies were produced in rabbits and affinity-purified using the recombinant antigen at ProteoGenix (Schiltigheim, France).

### Generation of Stwl knockout alleles

Stwl knockout (KO) alleles (replacement of protein coding sequence by a DsRed cassette) were generated using CRISPR-mediated homology directed repair. Briefly, 1000bp from the 3’UTR and 785bp from the 5’UTR of Stwl were cloned into a vector (pBSK-attB-DsRed-attB), flanked by a 3XP3-driven DsRed cassette. This plasmid was co-injected along with two gRNA-expressing plasmids (pU6-Bbs1-ChiRNA containing gRNA1: GATCCACTGGCTCTCGCTTA and gRNA2: GCATCAGGTTCACCTCAGAGG in embryos from the *nos-Cas9* strain (2^nd^ chromosome, BDSC78781) by Bestgene Inc. Transformants were selected based on DsRed expression and proper integration into the *stwl* locus was verified by PCR. Two independent and validated *stwl* KO alleles (*stwl^KO4^* and *stwl^KO7^*) were in our experiments.

### Fertility assays

For male fertility assays, two *yw* virgin females were crossed to a single tester male in a vial and allowed to mate for 1 week. Subsequently, the tester male was transferred to a new vial with two *yw* virgin females for the next week and so on. For each vial, the number of resulting progenies (F1) were counted until 20 days post setup. Female fertility assays were performed in a similar manner except that a single tester female was crossed to two ∼1d old *yw* males. Atleast 8 replicate crosses were set up for each genotype. Any vials that contained deceased parent flies were omitted from the analyses.

### Immunofluorescence staining and microscopy

For cultured cells, Kc167 cells were settled onto poly-L lysine coated glass slides at a concentration of 1×10^6^/ml for 2 hours. Cells were then fixed to the slide for 10 minutes with 4% formaldehyde in PBS-Triton (1x PBS with 0.01% Triton X-100) at room temperature and stored in PBS at 4°C until use. For the Stwl localization experiment, slides were instead fixed by methanol fixation. After settling cells onto slides for two hours as above, the slides were dipped into ice cold PBST (1x PBS with 0.02% Tween-20), incubated in cold methanol at −20°C for 10 minutes, and stored in PBS at 4°C until use. Cells were permeabilized in 1% Triton-PBS for 15 minutes and washed three times for 5 minutes each in PBST (1x PBS with 0.02% Tween-20). Slides were then blocked with BSA-PBST (1x PBS with 0.02% Tween-20 and 2% BSA) for 30 minutes with nutation. Primary antibodies were diluted in BSA-PBST, applied to the sample, and coverslips were sealed with rubber cement. Slides were incubated overnight at 4°C. The following day, slides were washed three times for 5 minutes each with PBST. Secondary antibodies were diluted in BSA-PBST, applied to samples, sealed with rubber cement, and incubated for 2 hours at room temperature while protected from light. Slides were washed three times for 5 minutes each with PBST. Slides were incubated with Hoescht (1:10,000 in 2xPBS) for 5 mins to stain DNA. Slides were then mounted using SlowFade Gold (Invitrogen).

For formaldehyde fixation and staining of *Drosophila* tissues, 3-4 ovaries or 5-7 testes per sample were dissected in 1XPBS and fixed in 4% EM-grade paraformaldehyde (PFA) for 20 min at room temperature (RT) on a nutator. Fixed samples were washed three times using 1xPBS containing 0.1% Triton-X (PBS-T) for 15 minutes each and blocked using 3% BSA in 1xPBS-T for 30 minutes. Primary antibodies were diluted in 3% BSA in 1xPBS-T block and added to the samples for overnight incubation at 4°C. On day two, samples were washed as above and incubated overnight at 4°C with secondary antibodies diluted in 3% BSA in 1xPBS-T. On day three, samples were washed as above and mounted with Vectashield + DAPI (Vector Laboratories). For methanol fixation and staining, 3-4 ovaries were dissected in 1xPBS and fixed in ice-cold 100% methanol for 10 min at −20°C. Following fixation, ovaries were washed and stained as above. The following primary antibodies were used in this study: rabbit anti-Stwl A2 (raised against full-length Stwl), mouse anti-Hts (1B1, 1:20, developmental studies hybridoma bank (DSHB)), rat anti-Vasa (1:100, DSHB), mouse anti-Lamin Dm0 (ADL84.12, 1:400; DSHB), mouse anti-Lamin C (LC28.26, 1:100; DSHB), mouse anti-Bam (1:50; DSHB), mouse anti-mAb414 (ab24609, 1:100; Abcam), rat anti-dCENP-A for Kc167 cells (AB_2793749, 1:100, Active motif), rabbit anti-dCENP-A for ovaries (AB_2793320, 1:200; Active Motif) and mouse anti-H3K9me2 (ab1220, 1:100, Abcam). Rabbit anti-Vasa (1:1000) was a gift from Prashanth Rangan. Guinea pig anti-Lamin Dm0, guinea pig anti-LBR and guinea pig anti-Otefin were gifts from Georg Krohne. All fluorescence microscopy images were acquired using a Leica TCS SP8 confocal microscope with 63x oil-immersion objectives (NA = 1.4). Z-stacks were acquired with a slice thickness of 0.30 μm for the FISH experiments and 0.50 μm for all other experiments.

### Immunofluorescence quantification and localization in Kc167 cells

IF images were analyzed using the ImageJ extension TANGO^73^. Stwl and lamin IF intensity was calculated for each nucleus using the integrated density function. For peripheral localization, images from methanol fixed IF samples were used. Nuclei were divided into 5 equi-volume shells using the shell analysis feature. The fraction of signal in the outer four shells were combined to create the peripheral compartment while the inner shell constituted the center compartment. The average peripheral to center ratio was calculated across three replicates.

### IF-Oligopaint DNA FISH

For *Drosophila* ovaries, whole mount tissue immunofluorescence was performed as mentioned above. Subsequently, samples were post-fixed with 4% PFA for 50 min and washed three times for 5 minutes each in 2xSSC containing 0.1% Tween-20 (2x SSC-T). Samples were then washed in 2xSSC-T with increasing formamide concentrations (20%, 40% and 50%) for 10 min each followed by a final 10 min wash in 50% formamide. Next, samples in 50% formamide + 2X SSC-T were transferred to a PCR tube and incubated at 37°C for 4 hr, 92°C for 3 min, and 60°C for 20 min. After this step, excess formamide solution was removed and the hybridization mix (20-40 pmol per probe, 36μl probe buffer + 1μl RNAse A) was added to the ovaries. Samples were denatured at 91°C for 3 min followed by overnight incubation at 37°C in the dark. Following hybridization, samples were first rinsed with 50% formamide + 2xSSC-T and then washed two times for 30 minutes each at 37°C. Next, samples were washed once with 20% formamide + 2xSSC-T for 10 min at RT followed by four washes with 2xSSC-T for 3 min each and then mounted with Vectashield + DAPI. Oligopaints targeting a 100kb region on Chr2R:23,799,747-23,900,018 were synthesized for *bgcn* locus DNA FISH. On a single slice, the shortest distance from the FISH focus to the nuclear periphery (marked by Vasa) was identified visually and measured using the line tool in the LAS X Leica software to estimate the NE-focus distances.

### RNA FISH

RNA FISH in ovaries was performed using the Stellaris RNA FISH protocol for imaginal discs with minor modifications. Briefly, 3-4 ovaries were dissected in ice cold RNase-free 1xPBS and fixed in 4% PFA in 1xPBS for 30 minutes on a nutator with gentle shaking. Following fixation, samples were washed three times with RNAse-free 1xPBS for 5 minutes each and incubated with 1ml 100% ethanol at 4°C overnight on a nutator. The next day, samples were washed with RNAse-free wash buffer A (2xSSC, 10% formamide) for 3 minutes at RT and incubated with 100l of hybridization mix (50-125nM probes, 2xSSC, 10% dextran sulfate, 1g/l *E.coli* tRNA, 2mM vanadyl ribonucleoside complex, 0.5% RNase free BSA, 10% deionized formamide, nuclease free water) overnight in a humid chamber at 37°C. Following the hybridization, the samples were washed twice with wash buffer A at 37°C for 30 minutes each, once wash buffer B for 5 min and mounted with Vectashield + DAPI. *bgcn* RNA FISH probes were designed using the Stellaris probe designer (Biosearch Technologies). polyT FISH probes were used to label mRNA and demarcate the nuclear boundary.

### Transmission electron microscopy

Ovaries were dissected and fixed in freshly prepared fixative (2.5 % glutaraldehyde in 0.1 M sodium cacodylate buffer). Fixed ovaries were stored at 4°C until sectioning. TEM was performed with the Center for Microscopy and Image Analysis at the University of Zürich. Image analysis was performed using Maps Viewer or ImageJ. Images were acquired such that each pixel corresponds to 1.7nm.

### RNA extraction from ovaries and RNA sequencing

Briefly, ovaries from 4–5-day old females were dissected in RNase-free 1X PBS and flash frozen in liquid nitrogen until RNA extraction. RNA extraction for each replicate was performed using 35 ovaries, using the RNeasy RNA extraction kit (Qiagen). Samples were treated with DNase post RNA extraction and purified using an RNA purification kit (Promega). RNA concentrations were assessed using a Nanodrop as well as a Qubit RNA analyzer for sample quality and RIN scores. Samples of sufficient quality (RIN>9) were subjected to library preparation (Illumina Truseq mRNA kit) followed by sequencing using Illumina Novaseq 6000 (single read, 100bp) at the Functional Genomics Center Zürich (FGCZ).

### RNA sequencing data analysis

On average, we generated 28.3 million reads per sample. The resulting raw reads were cleaned by removing adaptor sequences, low-quality-end trimming and removal of low-quality reads using BBTools v 38.18 [Bushnell, B. *BBMap*. Available from: https://sourceforge.net/projects/bbmap/.]. The exact commands used for quality control can be found on the Methods in Microbiomics webpage [Sunagawa, S. *Data Preprocessing — Methods in Microbiomics 0.0.1 documentation*. https://methods-in-microbiomics.readthedocs.io/en/latest/preprocessing/preprocessing.html]. Transcript abundances were quantified using Salmon v 1.10.1^74^ and BDGP6.32. Differential gene expression analysis was performed using Bioconductor R package DESeq2 v1.37.4^75^.

### Stwl CUT & RUN

CUT&RUN was performed as described in Kotb and colleagues^21^. Briefly, 20 pairs of fly ovaries were dissected per replicate and placed on ice in 1X PBS. Each sample was then treated with the permeabilization buffer (50mL PBST 500 μL Triton-X) for 1 hour at RT while nutating, followed by washing with 1 mL BBT+ buffer (0.5 g BSA final 0.5% 50 ml PBST) and subsequent removal of the supernatant. Antibody dilutions were prepared in 500 μL BBT+ buffer, and the sample was incubated overnight at 4°C. Next day, the sample was washed with PBT+ buffer and then incubated with pAG-MNase (1:100) in 500 μL BBT+ for 4 hours at room temperature. For DNA cleavage, the samples were resuspended in 150 μL Wash+ buffer (20mM HEPES, pH 7.5, 150mM NaCL, 0.1 % BSA, Roche complete EDTA-free tablet +0.5 mM spermidine) and incubated for 45 minutes at 4°C. The reaction was stopped by adding 150 μL 2xSTOP buffer (200 mM NaCl, 20 mM EDTA) for 30 minutes at 37°C. The sample was then centrifuged at 16,000g for 5 minutes and the supernatant was carefully extracted and transferred to a fresh eppendorf tube. 2 μL of 10% SDS and 2.5 μL of 20 mg/mL Proteinase K was added to the supernatant and the mixture was thoroughly mixed using a brief vortexing procedure. Subsequently, the sample was incubated at 50°C in a water bath for 2 hours. It’s important to note that this can be stopped at this step and the samples can be stored at −-20 C. 20 μL of AmpureXP bead slurry and 280 μL of MXP buffer (20% PEG8000, 2.5 M NaCl, 10 mM MgCl2) were added to 150 μL of the supernatant and incubated for 15 minutes at RT. Using a magnetic rack, the beads were collected and the supernatant was discarded. While on the magnetic rack, 1 mL of 80% ethanol was added to each tube without disturbing the beads. The sample was then incubated for a minimum of 30 seconds and the ethanol was gently aspirated until all traces of ethanol were removed. The beads were then air-dried for 2 minutes, resuspended in 10 μL of RNAse-free and DNAse-free water and incubated at RT for 2 minutes. The clear solution (containing the liberated DNA) was then transferred to a new eppendorf tube. The DNA concentration was determined using a dsDNA high-sensitive Qubit assay and analyzed DNA size distribution in samples using a Fragment analyzer.

### CUT & RUN library preparation and data analysis

The NEBNext Ultra II DNA Library Prep Kit for Illumina (E7645, E7103) protocol was followed for library preparation. Reads were first evaluated for their quality using FastQC (v0.11.8, RRID:SCR_014583). Reads were trimmed for adaptor sequences using Trim Galore! (v0.6.6, RRID:SCR_011847) and aligned to the dm6 reference genome version for *Drosophila melanogaster* using Bowtie2 (version 2.2.8 RRID:SCR_016368) with parameters -q -I 50 -X 700 --very-sensitive-local --local --no-mixed --no-unal --no-discordant. Binary alignment maps (BAM) files were generated with samtools v1.9 and were used in downstream analysis. MACS2 v2.1.0 was used to call significant peaks for samples. IgG was used as control to call peaks. Peaks within ENCODE blacklisted regions and repetitive sequences larger than 100 bases were removed. Coverage tracks were generated from BAM files using deepTools 3.2.1 bamCoverage function with parameters– normalize using RPKM–bin size 10. For genomic annotation promoters (−500 b to +500 b) relative to the TSS were defined according to the drosophila dm6 reference genome version. ChipSeeker (v1.36.0) was used to annotate Stonewall peaks. Heatmaps of genomic regions were generated with deepTools 3.2.1 computeMatrix and plotHeatmap commands, or EnrichedHeatmap (v1.30.0). PCA plot of histone modifications was generated using deepTools 3.2.1 multiBigwigSummary and plotPCA functions.

### Affinity purification and mass spectrometry

Approximately 1.5 x 10^8^ Kc167 cells were harvested for each replicate and stored at −80°C until further use. For lysis, cells were thawed and resuspended in a buffer containing 50mM Tris HCl (pH 7.4), 150mM NaCl, 0.3mM MgCl_2_, 5% glycerol, 0.5% NP40, protease inhibitor cocktail (PIC), 1X PMSF and Benzonase. Lysis was performed using 25 strokes of a type B pestle followed by a one-hour incubation at 4°C. Lysates were centrifuged at 4300g for 25 minutes at 4°C and the resulting supernatant was transferred into a fresh tube. Protein concentration was estimated using BCA method. For the affinity purification, lysates with equal protein concentration were incubated with Rabbit IgG (Merck, control) and 50 μg of Stwl antibody overnight at 4°C. Next day, pre-equilibrated magnetic Protein A/G beads (125μl slurry/ sample) were added to each sample at room temperature for ∼1.5 hours while rotating. Following this, beads were washed once with lysis buffer and twice with bead wash buffer (50mM Tris HCl pH 7.4 and 150mM NaCl). Washed beads with bound protein complexes were subjected to proteolysis by on-bead digestion. Samples were transferred into a 10 kDa molecular weight cutoff spin column (Vivacon 500, Sartorious), following the FASP protocol^76^. Beads in solution were dried, denaturated (8M Urea), reduced (5mM TCEP, 30min 37°C) and alkylated (10mM Iodoacetamide, 30min 37°C). Beads were then washed three times with 50mM ammonium bicarbonate (250µl). During the buffer exchange, samples were centrifuged at 10000g. Subsequently, samples were proteolyzed with 0.5µg of Trypsin (Promega, sequencing grade) for 16h at 37°C. The proteolysis was quenched with 5% formic acid and peptides were subjected to C18 cleanup (BioPureSPN PROTO 300 C18, Nest group), following the manufacturer’s procedure. The eluted peptides were then dried using a speedvac and resuspended in 20µl of 2% acetonitrile and 0.1% formic acid. LC-MS/MS was performed on an Orbitrap Exploris 480 mass spectrometer (Thermo Fisher) coupled to an Vanquish Neo liquid chromatography system (Thermo Fisher). Peptides were separated using a reverse phase column (75 μm ID x 400 mm New Objective, in-house packed with ReproSil Gold 120 C18, 1.9 μm, Dr. Maisch GmbH) across 180 min linear gradient from 7 to 50% (buffer A: 0.1% [v/v] formic acid; buffer B: 0.1% [v/v] formic acid, 80% [v/v] acetonitrile). Samples were acquired in DDA mode (Data Dependent Acquision) with MS1 scan (scan range = 350-1500, R=60K, max injection time auto and AGC target = 100), followed by 30 dependent MS2 scans (scan range = 120-2100, R = 30K, max injection time auto and AGC target = 200). Peptides with charge between 2-6 were isolated (m/z = 1.4) and fragmented (NCE 28%). Acquired spectra were analyzed using the MaxQuant software version 1.5.2.8 against the *Drosophila* proteome reference dataset (http://www.uniprot.org/, downloaded on 18.01.2021, 22’044 proteins including not reviewed proteins) extended with reverse decoy sequences. The search parameters were set to include specific tryptic peptides, maximum two missed cleavage, carbamidomethyl as static peptide modification, oxidation (M) and deamidation (N-terminal) as variable modification and “match between runs” option. The MS and MS/MS mass tolerance was set to 10 ppm. False discovery rate of < 1% was used at PSM and protein level. Protein abundance was determined from the intensity of top two unique peptides. Intensity values of proteins identified in all replicates in at least one condition (Stwl pulldown or control pulldown) were median normalized and imputed using random sampling from a normal distribution generated from 1% lower values. Statistical analysis was performed using unpaired two-sided t-test. Hits identified from the differential analysis between the Stwl pulldown versus the IgG control, with log_2_FC>1 and p-value<0.05, were considered as interacting proteins.

### Egg chamber classification and quantification

Ovaries from *bam-Gal4 > mCherry^RNAi^*or *bam-Gal4 > Stwl^RNAi^*females were dissected in 1xPBS followed by the addition of Vectashield containing DAPI. Ovarioles were gently separated and mounted on a glass slide. Egg chamber stages were classified and quantified as described elsewhere^77^.

## Acknowledgements

We are indebted to the members of the Jagannathan lab, Joyce lab, Rangan lab, Gabriel Neurohr and Tatjana Kleele for their comments on the manuscript. We thank Hugo Stocker for his valuable suggestions throughout the course of the project. We are grateful to Dan Hasson from the BiNGS core for assistance with the CUT&RUN data analysis and Shinichi Sunagawa from ETH Zürich for providing bioinformatics resources. We thank the BDSC, VDRC, BDGP Gene Disruption Project, and Flybase for reagents and resources. We thank Georg Krohne and Victor Corces for sharing antibodies. We thank the Functional Genomics Center Zurich (FGCZ), Center for Microscopy and Image Analysis and, Scientific Center for optical and Electron Microscopy (ScopeM), the Mass Spectrometry facility at the IBC ETH Zurich for technical support. M.J is supported by a project grant (310030_189131) from the Swiss National Science Foundation. P.R. is funded by grants from the NIH/NIGMS (R01GM111779, RO1GM135628 and R56AG082906). E.F.J is funded by grants from the NIH/NIGMS (R35GM128903), NIH/NICHD (R21HD107261) and the NSF (2207050). U.K. is funded by a project grant (310030_219203) from the Swiss National Science Foundation. A.C acknowledges support from Genetics Society of America in the form of a DeLill Nasser Award for Professional Development in Genetics. This work was supported in part by the Bioinformatics for Next Generation Sequencing (BiNGS) shared resource facility within the Tisch Cancer Institute at the Icahn School of Medicine at Mount Sinai, which is partially supported by NIH grant P30CA196521. This work was also supported in part through the computational resources and staff expertise provided by Scientific Computing at the Icahn School of Medicine at Mount Sinai and supported by the Clinical and Translational Science Awards (CTSA) grant UL1TR004419 from the National Center for Advancing Translational Sciences. Research reported in this paper was also supported by the Office of Research Infrastructure of the National Institutes of Health under award number S10OD026880.

## Author Contributions

E.F.J. and M.J., A.C. and R.I. conceived the project. A.C., R.I., S.C.N., N.K., J.H. designed and performed most of the experiments, except for Stwl purification performed by C.A. G.U. performed the CUT&RUN data analysis and A.S. performed the RNA-seq data analysis. AP-MS was performed with the help of F.U., who also analyzed the data. E.F.J, M.J, A.C. and R.I. wrote the manuscript with input from all authors.

## Declaration of interests

The authors declare no competing interests.

---

**Figure S1.**
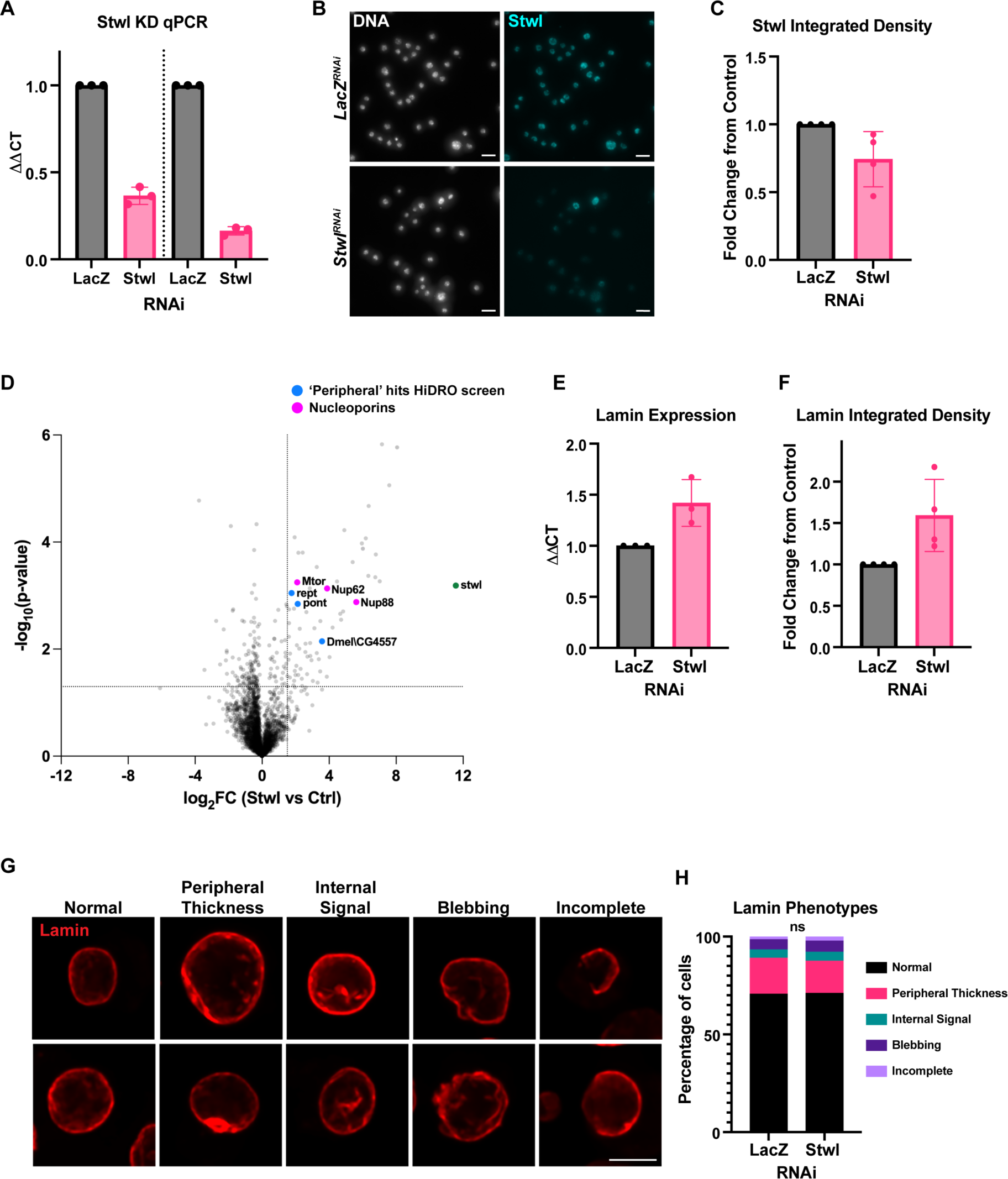
(A) qPCR for Stwl following LacZ RNAi (control) and Stwl RNAi treatment. The ΔΔCT was calculated using two different Stwl qPCR primers across three replicates. (B) Immunofluorescence against Stwl (blue) in control (lacZ RNAi) or Stwl RNAi treated Kc167 cells stained for DNA (grey). (C) Change in integrated density of Stwl immunofluorescence signal from Kc167 cells across four replicates. Each dot represents the fold change between medians of one replicate. Each replicate contained >300 nuclei. (D) Volcano plot of the Stwl-associated proteome in Kc167 cells from three biological replicates. The dashed lines mark log_2_FC>1.5 and p<0.05; magenta points indicate NPC-associated nucleoporins and blue points indicate proteins identified as peripheral hits from the HiDRO screen. (E) qPCR for Lamin B following LacZ RNAi (control) and Stwl RNAi treatment. The ΔΔCT was calculated across three replicates. (F) Change in integrated density of Lamin B immunofluorescence signal from Kc167 cells across four replicates. Each dot represents the fold change between medians of one replicate. Each replicate contained >300 nuclei. (G) Categorizations of Lamin B phenotypes. Two example nuclei are shown for each phenotype. (H) Quantification of lamin phenotypes in control (LacZ RNAi) and Stwl RNAi Kc167 cells from three replicates. No significant changes were found using a Chi-square test.

**Figure S2.**
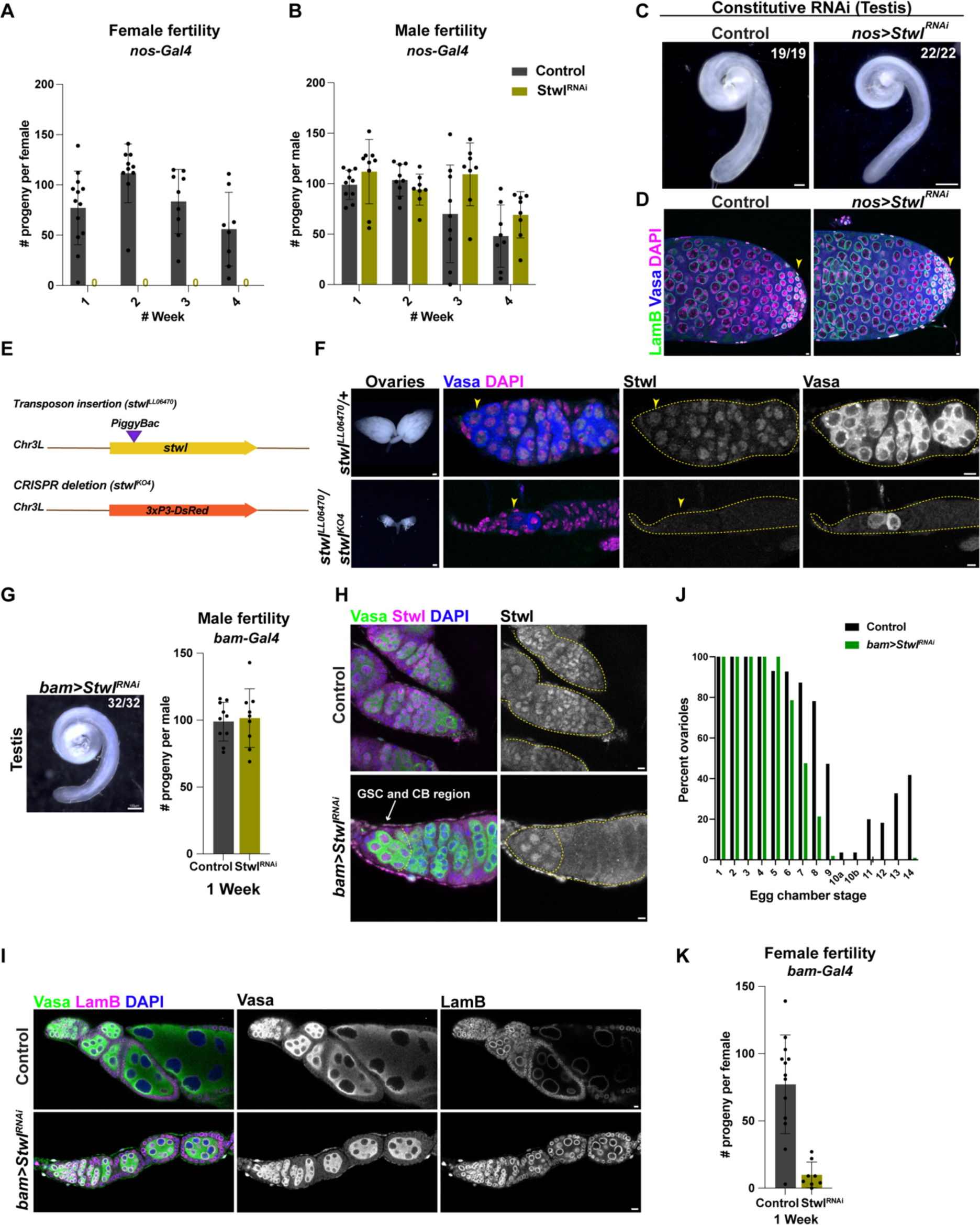
Stwl is required for GSC maintenance and female fertility. (A, B) Fertility assay of females (A) and males (B) from *TM3* / *Stwl^RNAi^* (Control) and *nos > Stwl^RNAi^* flies over four weeks. Each dot represents number of progenies sired by a single female (A) or male (B) fly from n≥8 crosses. (C) Testes from *TM3 / Stwl^RNAi^* (Control) and *nos > Stwl^RNAi^* males at 3 days post eclosion. Scale bar:100μm. (D) Apical tip of testes from *TM3 / Stwl^RNAi^* (Control) and *nos* > *Stwl^RNAi^* males stained for Lamin B (green), Vasa (blue) and DAPI (magenta). Scale bar:5μm. (E) *stwl* mutant alleles used in this study – a *PiggyBac* insertion in the Stwl locus (*stwl^LL6470^*) and a CRISPR/Cas9-mediated knockout of the Stwl coding sequence (*stwl^KO4^*). (F) Ovaries (left panel) and germaria (right panel) from *stwl^LL6470^*/+ (Control) and *stwl^LL6470^*/*stwl^KO4^* females 3d post eclosion stained for Vasa (blue) and Stwl (magenta). Scale bar (ovaries):100μm. Scale bar (germaria):5μm. (G) Testis (left panel) and male fertility assay (right panel) from *bam > mCherry^RNAi^*(Control) and *bam > Stwl^RNAi^* males at 3 days post eclosion. Each dot in the right panel represents number of progenies sired by a single male from n≥9 crosses. Scale bar:100μm. (H) Germaria from *bam > mCherry^RNAi^* (Control) and *bam > Stwl^RNAi^* ovaries stained for Stwl (magenta), Vasa (green) and DAPI (blue). Scale bar:5μm. (I) Ovarioles from *bam > mCherry^RNAi^* (Control) and *bam > Stwl^RNAi^* ovaries stained for Lamin B (magenta), Vasa (green) and DAPI (blue). Scale bar:5μm. (J) Quantification of egg chamber stages from (I). n=55 ovarioles from *bam > mCherry^RNAi^* (Control) and n=103 ovarioles from *bam* > *Stwl^RNAi^*. (K) Fertility assay of females from *bam > mCherry^RNAi^* (Control) and *bam > Stwl^RNAi^*over one week. Each dot represents number of progenies sired by a single female from n≥8 crosses.

**Figure S3.**
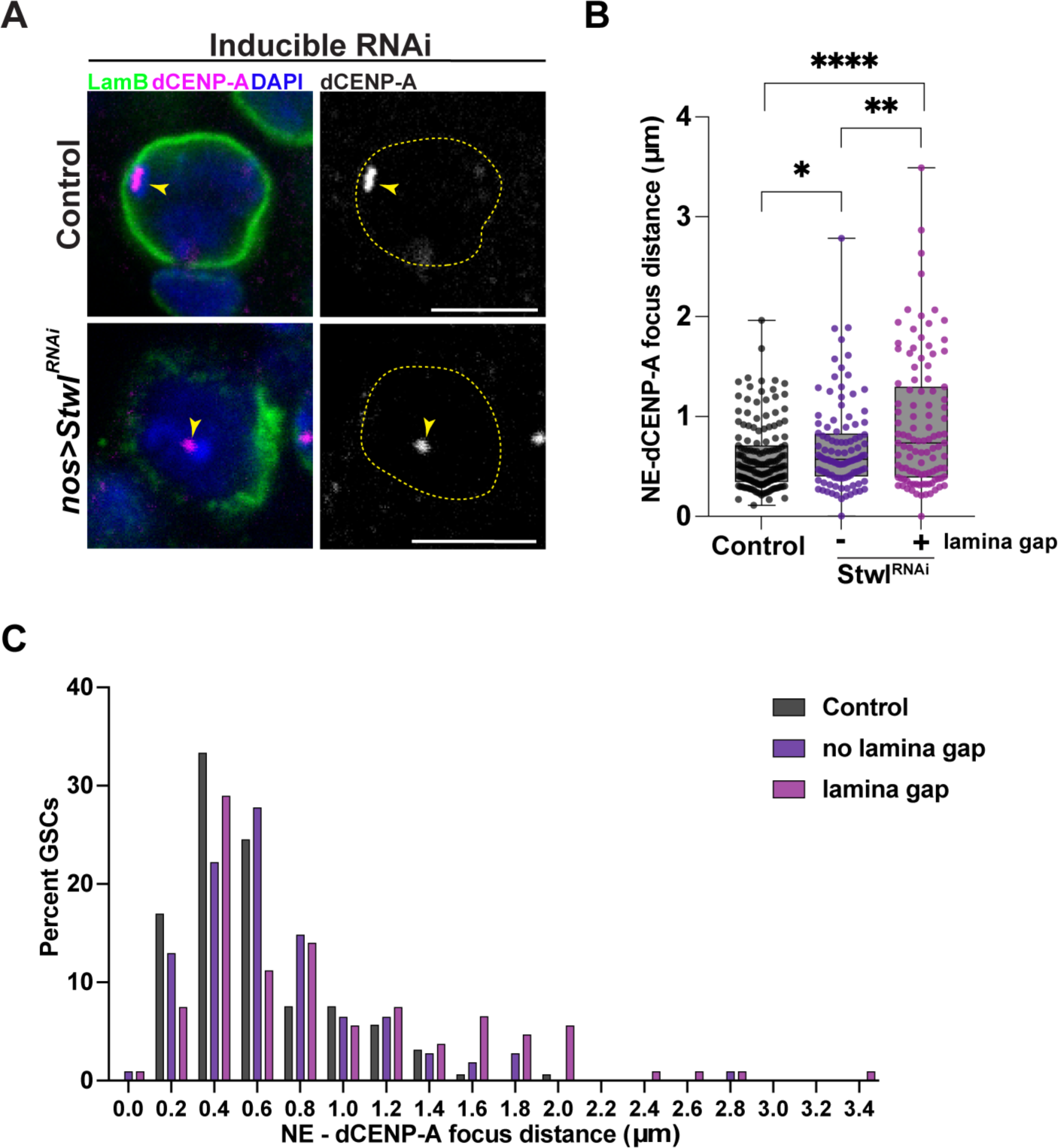
Loss of Stwl leads to a reduction of NE proximal centromeres. (A) IF staining of dCENP-A (magenta), Lamin B (green) and DAPI (blue) in GSCs from *nos > mCherry^RNAi^* (Control) and *nos* > *Stwl^RNAi^* ovaries following a 4d shift to 29°C. Yellow arrowheads point to centromeric foci and yellow dotted lines indicate the nuclear boundary. Scale bar:5μm. (B) Quantification of NE-centromere distance (µm) in GSCs from Control (n=159 foci) and *nos* > *Stwl^RNAi^* (n= 215 foci). * indicates p<0.05, ** indicates p<0.01 and **** indicates p<0.0001 from Student’s t-test. (C) Histogram of NE-centromere distance (µm) in GSCs from (B).

**Figure S4:**
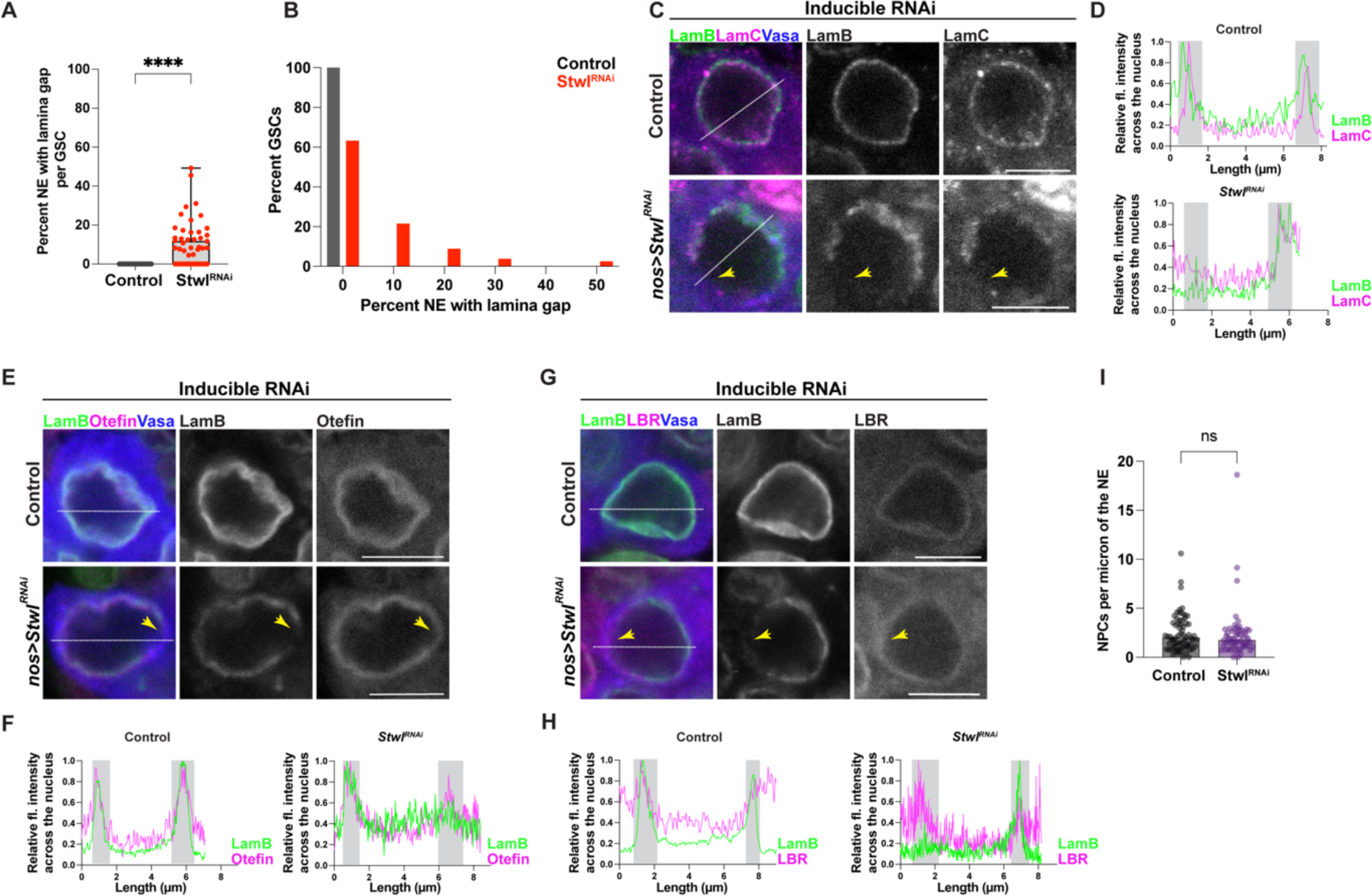
Stwl knockdown in female GSCs leads to substantial changes at the nuclear envelope. (A) Quantification of the percentage of the NE with a lamina gap in GSCs from *nos > mCherry^RNAi^* (Control; n=78) and *nos > Stwl^RNAi^* (n=78) ovaries following a 4d shift to 29°C. **** indicates p<0.0001 from Student’s t-test. (B) Histogram showing the percentage of GSCs from (A) with lamina gaps. (C) IF staining of Lamin B (green), Lamin C (magenta) and Vasa (blue) in GSCs from *nos > mCherry^RNAi^* (Control) and *nos > Stwl^RNAi^*ovaries following a 4d shift to 29°C. Scale bar:5μm. (D) Relative fluorescence intensity of Lamin B (green) and Lamin C (magenta) across the nucleus (white dotted line) from (C). Shaded grey regions mark the NE. (E) IF staining of Lamin B (green), Otefin (magenta) and Vasa (blue) in GSCs from *nos > mCherry^RNAi^* (Control) and *nos > Stwl^RNAi^* ovaries following a 4d shift to 29°C. Yellow arrowheads show lamin gaps. Scale bar:5μm. (F) Relative fluorescence intensity of Lamin B (green) and Otefin (magenta) across the nucleus (white dotted line) from (E). Shaded grey regions mark the NE. (G) IF staining of Lamin B (green), LBR (magenta) and Vasa (blue) in GSCs from *nos > mCherry^RNAi^* (Control) and *nos > Stwl^RNAi^* ovaries following a 4d shift to 29°C. Yellow arrowheads show lamin gaps. Scale bar:5μm. (H) Relative fluorescence intensity of Lamin B (green) and LBR (magenta) across the nucleus (white dotted line) from (G). Shaded grey regions mark the NE. (I) Number of NPCs per micron of the NE were quantified from TEM images of GSC-like cells from *nos > mCherry^RNAi^* (Control; n=66) and *nos > Stwl^RNAi^* (n=59) ovaries in a *bam^Δ86^/bam^1^* background following a 4d shift to 29°C. ns indicates p>0.05 from Student’s t-test.

**Figure S5.**
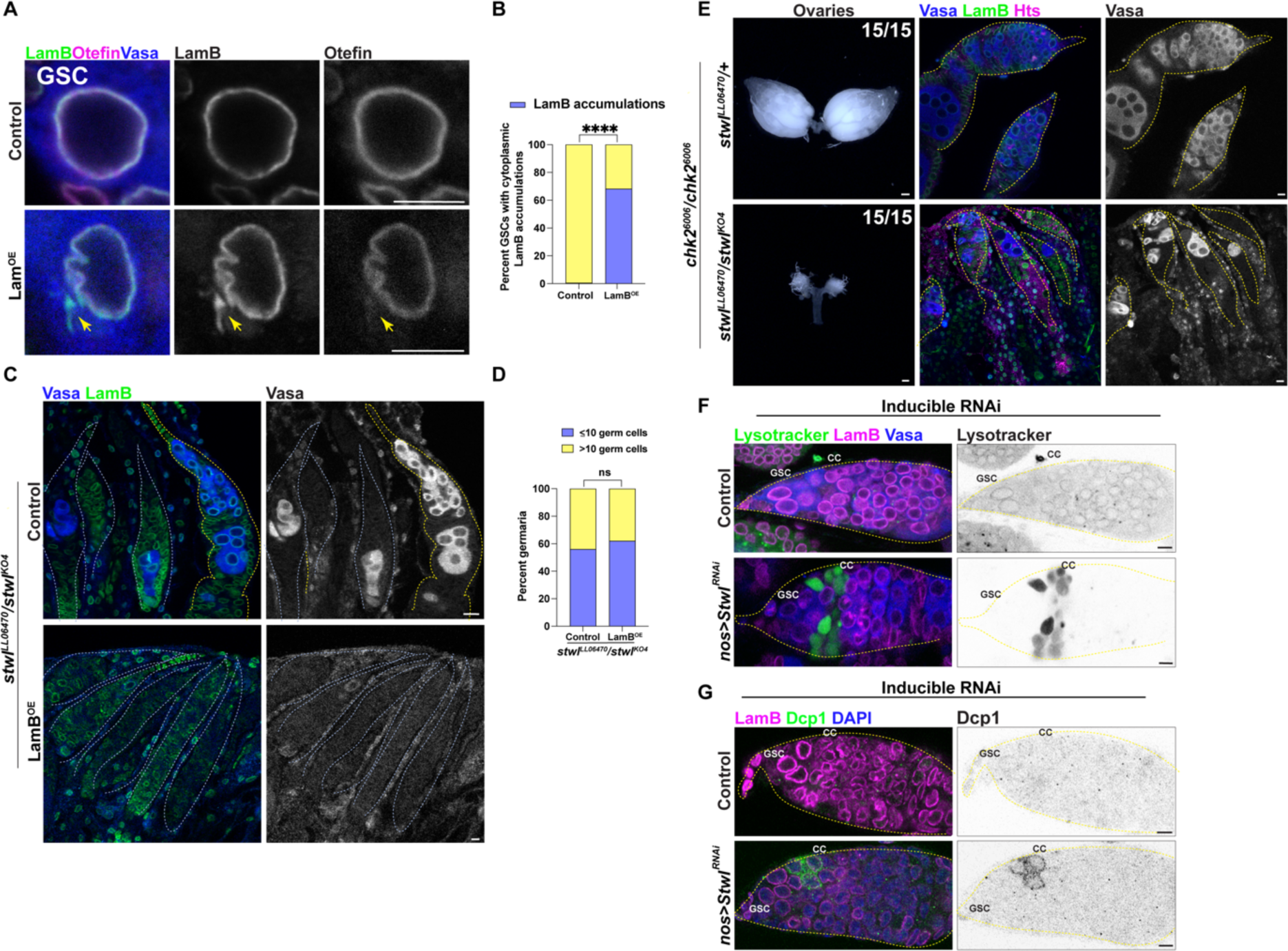
GSC loss upon Stwl knockdown is not dependent on Lamin B levels or Chk2 signaling. (A) IF staining of Lamin B (green), Otefin (magenta) and Vasa (blue) in GSCs from *nos*/*TM3* (Control) and *nos > Lam^EY08333^* (Lam^OE^) ovaries. Yellow arrowheads indicate cytoplasmic lamin B accumulations in lamin overexpressing GSCs. Scale bar:5μm. **** indicates p>0.0001 from Student’s t-test. (B) Percentage of GSCs from (A) with cytoplasmic lamin accumulations. n=188 GSCs from the control and n=176 GSCs from *Lam^OE^*. **** indicates p<0.0001 from Fisher’s exact test. (C) IF staining of Lamin B (green) and Vasa (blue) in germaria from control and *nos > Lam^EY08333^* (Lam^OE^) ovaries in a *stwl^KO4^/stwl^LL6470^* background. Scale bar:5μm. (D) Percent of germaria with the indicated number of Vasa positive germ cells from (C). n=75 ovarioles from the control and n=58 ovarioles from *Lam^OE^*. **** indicates p<0.0001 and ns indicates p>0.05 from sswtudent’s t-test. (E) Ovaries (left panel) and germaria (right panel) from *stwl^0LL6470^*/+ (Control) and *stwl^LL6470^*/*stwl^KO4^* females ovaries in a *chk2*^6006^ background stained for Lamin B (green), Hts (magenta) and Vasa (blue). Yellow dotted lines indicate the germarium boundary. Scale bar (ovaries):100µm. Scale bar (germaria):5 μm. (F) IF staining of Lysotracker (green), Lamin B (magenta) and Vasa (blue) in germaria from *nos > mCherry^RNAi^* (Control) and *nos > Stwl^RNAi^* following a 4d shift to 29°C. CC refers to germline cyst cells. Scale bar:5μm. (G) IF staining of Dcp-1 (green), Lamin B (magenta) and Vasa (blue) in germaria from *nos > mCherry^RNAi^* (Control) and *nos > Stwl^RNAi^* following a 4d shift to 29°C. CC refers to germline cyst cells. Scale bar:5μm.

**Figure S6.**
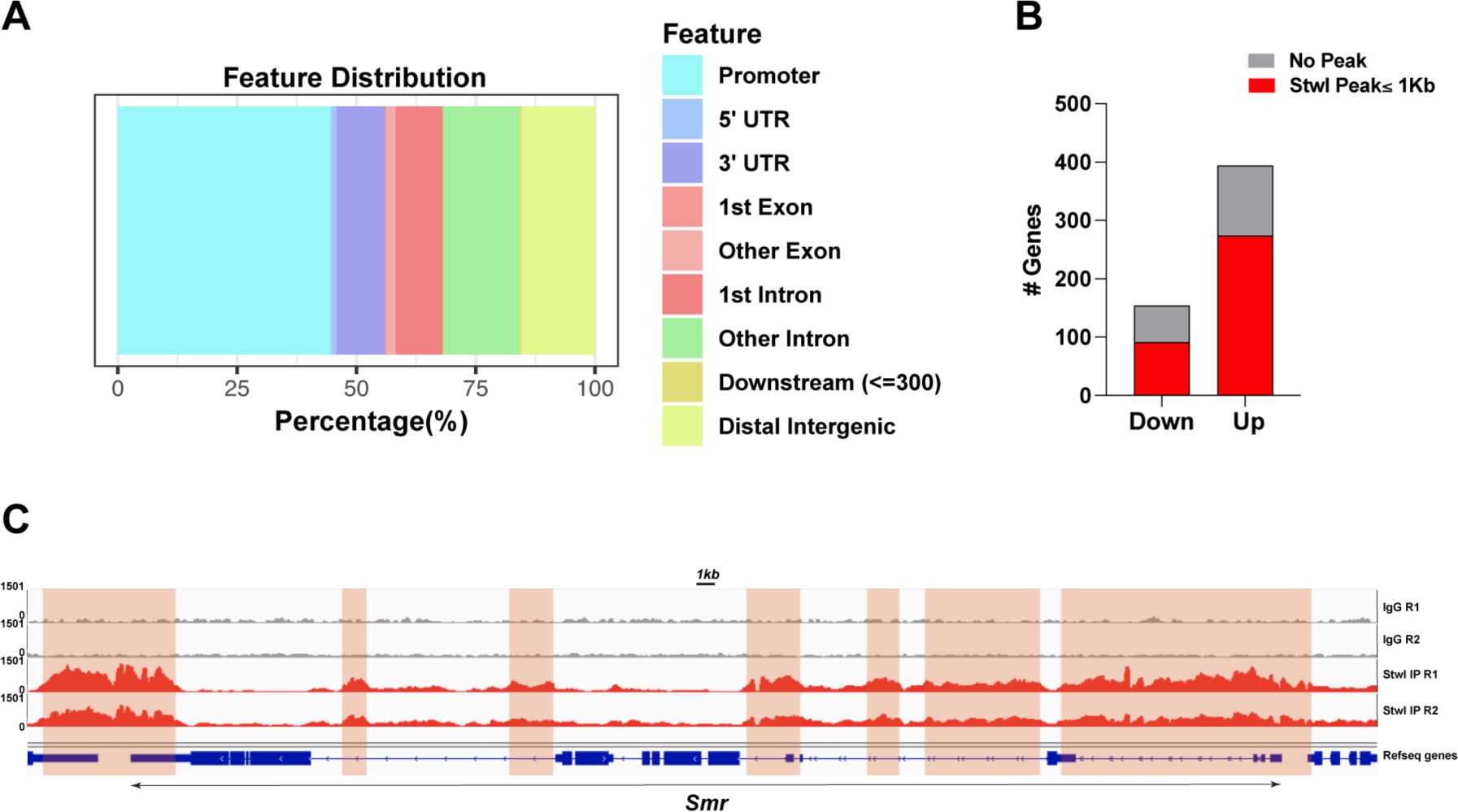
Stwl binds at the promoters and UTRs of regulated genes. (A) Percentages of Stwl CUT&RUN binding peaks at the indicated genomic regions. (B) Number of downregulated and upregulated genes with a Stwl peak within 1kb of the gene body following Stwl^OE^ in GSC-enriched ovaries. (C) Capture of the IGV genome browser (v2.11.4) showing an approximately 70kb region on the *Drosophila* X chromosome (y axis = reads per kilobase per million reads). Ensembl genes (blue). Shaded areas correspond to Stwl binding peaks.

## Table Legends

**Table S1. z scores for the indicated paramters from the HiDRO screen.**

**Table S2. List of proteins detected by LC-MS/MS in Kc167 lysates from control and Stwl affinity purification.**

**Table S3. Differentially expressed genes following Stwl overexpression in ovaries enriched for GSC-like cells.**

**Table S4. Normalized reads following Stwl overexpression in ovaries enriched for GSC-like cells.**

## Notes

### Competing Interest Statement

The authors have declared no competing interest.

